# AlphaFold2 structures template ligand discovery

**DOI:** 10.1101/2023.12.20.572662

**Authors:** Jiankun Lyu, Nicholas Kapolka, Ryan Gumpper, Assaf Alon, Liang Wang, Manish K. Jain, Ximena Barros-Álvarez, Kensuke Sakamoto, Yoojoong Kim, Jeffrey DiBerto, Kuglae Kim, Tia A. Tummino, Sijie Huang, John J. Irwin, Olga O. Tarkhanova, Yurii Moroz, Georgios Skiniotis, Andrew C. Kruse, Brian K. Shoichet, Bryan L. Roth

## Abstract

AlphaFold2 (AF2) and RosettaFold have greatly expanded the number of structures available for structure-based ligand discovery, even though retrospective studies have cast doubt on their direct usefulness for that goal. Here, we tested unrefined AF2 models *prospectively*, comparing experimental hit-rates and affinities from large library docking against AF2 models vs the same screens targeting experimental structures of the same receptors. In *retrospective* docking screens against the σ_2_ and the 5-HT2A receptors, the AF2 structures struggled to recapitulate ligands that we had previously found docking against the receptors’ experimental structures, consistent with published results. *Prospective* large library docking against the AF2 models, however, yielded similar hit rates for both receptors versus docking against experimentally-derived structures; hundreds of molecules were prioritized and tested against each model and each structure of each receptor. The success of the AF2 models was achieved despite differences in orthosteric pocket residue conformations for both targets versus the experimental structures. Intriguingly, against the 5-HT2A receptor the most potent, subtype-selective agonists were discovered via docking against the AF2 model, not the experimental structure. To understand this from a molecular perspective, a cryoEM structure was determined for one of the more potent and selective ligands to emerge from docking against the AF2 model of the 5-HT2A receptor. Our findings suggest that AF2 models may sample conformations that are relevant for ligand discovery, much extending the domain of applicability of structure-based ligand discovery.

## Introduction

Structure-based library docking is widely used in early ligand discovery^1^; on targets with well-formed binding sites^2,3^, the affinities of direct docking hits can reach the mid-nanomolar or even high picomolar range^4–13^. These docking campaigns have mostly relied on experimental protein structures from crystallography or, more recently, cryoEM, but for many drug targets^14^, such experimental structures remain unavailable. In such cases, template-based homology models have been used^15–17^. These models show promise, particularly when the sequence identity between the target and template exceeds 50%^18^. Many targets are outside of this range, and docking against homology models is often thought to reduce performance relative to experimental structures.

Recent breakthroughs in protein structure predictions, led by deep-learning methods such as AF2^19^ and RosettaFold^20^, promise to overcome this limitation. AF2 has demonstrated an unprecedented ability to predict protein structures with atomic accuracy^19^ and operates at scale^21,22^. As of this writing, the AlphaFold database features protein structures for the human proteome and for the proteomes of 47 other key organisms, covering over 200 million proteins and nearly all potential therapeutic protein targets. These structures have proven remarkably useful for multiple applications including structural biology^23–25^, protein design^26,27^, protein-protein interaction^28–30^, target prediction^31,32^, protein function prediction^33,34^, and biological mechanism of action^35–37^.

The impact of AF2 structures on structure-based ligand discovery has been murkier. While the global accuracy of the models has been impressive, concerns have arisen regarding their accuracy in modeling ligand binding sites^38^, where high fidelity to low energy conformations is thought to be crucial, and small errors, relatively unimportant for other applications, can disrupt ligand recognition and pose prediction. Indeed, retrospective studies have suggested that unrefined AF2 models struggle to recognize and pose known ligands, compared to decoy molecules, versus experimental structures in apples-to-apples comparisons ^7,39–48^. A drawback to these studies is that they are biased by the past; that is, known ligands can affect the conformations adopted by experimental structures when they are determined in complex with them, while experimental structures can influence the exploration of new ligands. Thus, it is possible that the relatively poor performance of AF2 models in *retrospective simulation* of structure-based ligand discovery may underestimate the ability of AF2 structures to template new ligand discovery *prospectively*.

Here, we begin to address this prospective gap. We selected two targets where the AF2 models appeared before the crystal structures were released, reducing possible bias: the σ_2_ receptor and the 5-HT2A serotonin receptor, belonging to two unrelated protein families, the EXPERA family and the G Protein Coupled Receptor (GPCR) family, respectively. In both models, AF2 predicted uncollapsed binding sites, unlike several other targets where we judged the orthosteric sites in the AF2 models too compressed to support plausible ligand fitting (**Extend Data Fig. 1**). For σ_2_, the AF2 structure recapitulated the side chain conformations of all orthosteric site residues to 1.1 Å RMSD versus the crystal structure, with few residues with individual RMSD values greater than 1.5 Å (**Figure 1**). For the 5-HT2A receptor, while most binding site residues were well predicted (RMSD < 2 Å versus the cryo-EM structure), two residues differed by 2.5 to 3.1 Å RMSD, adopting different rotamers (**Figure 1**). Thus, the σ_2_ and 5-HT2A receptor occupy two of what are perhaps three categories of AF2 models for ligand discovery applications: (1) those that are close to the experimental ligand binding site, though still harboring meaningful side chain differences: (2) those that are overall close but have several residues in substantially different conformations; and a third class where the modeled sites differ so much from the experimental as to preclude reliable ligand recognition without extensive refinement (**Extended Data Fig. 1**). We set out to run identical large library docking screens against the AF2 models and the experimental structures for both the σ_2_ and 5-HT2A receptors (i.e., four prospective large library docking campaigns overall), representing the first two classes. From each of these four campaigns we prioritized new chemotypes for synthesis and testing based on docking scores against each structure of each target (one from AF2, one experimental for each receptor). To ensure that the comparisons were meaningful, we synthesized and tested hundreds of predicted molecules against each receptor model and structure. We assessed the performance of the AF2 versus the experimental structures by hit rate (number experimentally active/number tested), chemotype diversity of the top-ranking list, and by the potency distribution of experimental actives. Our results suggest that AF2 structures may be more relevant to *prospective* structure-based ligand discovery than thought based on retrospective simulation.

**Figure 1.**
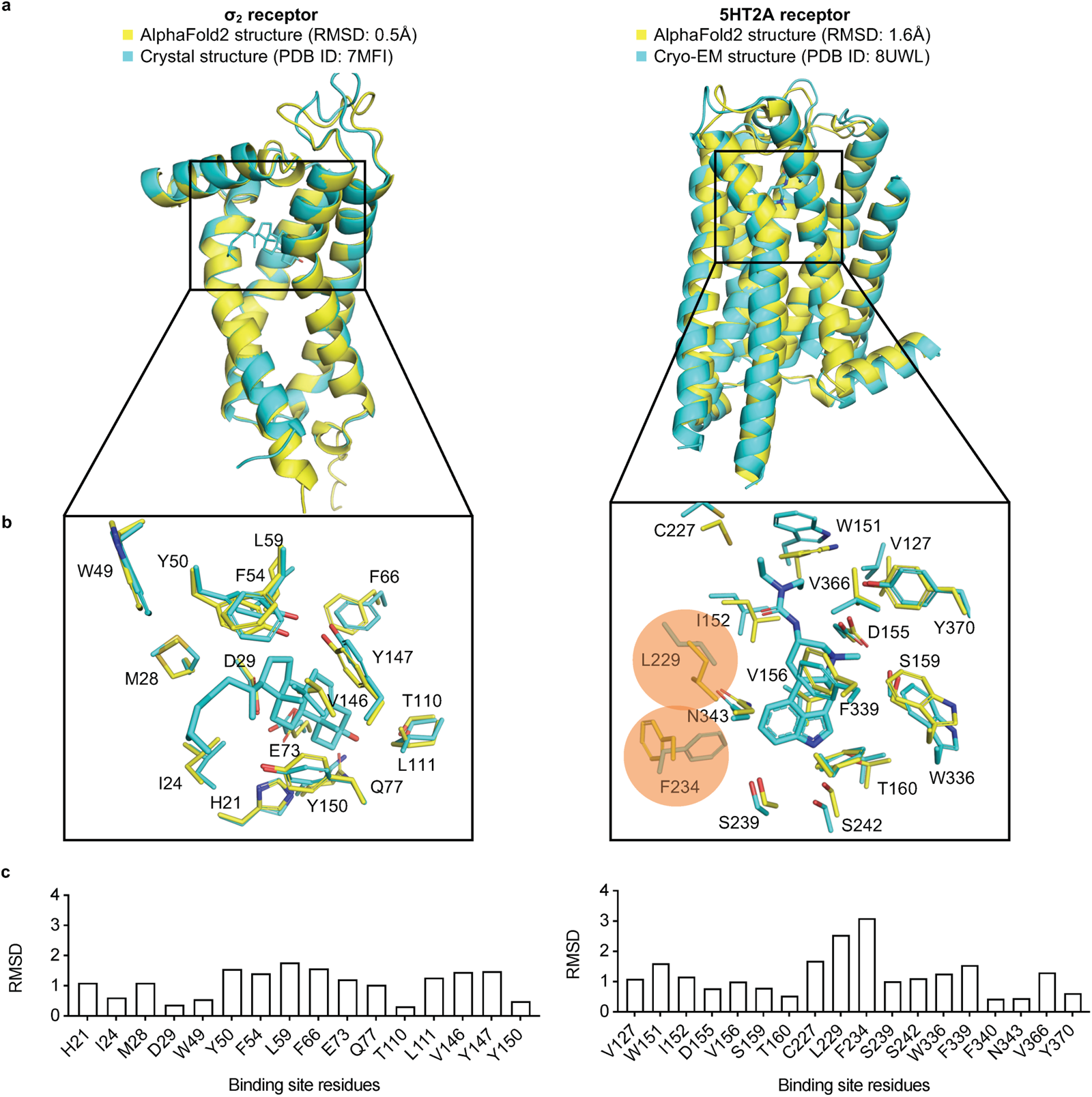
Structural comparisons of the σ_2_ receptor (left column) and the 5-HT2A receptor (right column) between the AlphaFold2 (AF2) predicted structure and the experimental structure. **a**. The experimental structure (in cyan) is overlaid with the AF2 predicted structure (in yellow). The Root Mean Square Deviation (RMSD) value is calculated based on backbone atoms. The ligand binding site residues were selected within 5 Å distance from the ligand. **b**. The full-atom RMSD values of the binding site residues between the AF2 and the experimental structures. Two residues with large conformational differences between the AF2 and experimental structures used in docking, Leu229 and Phe234, are highlighted for the 5-HT2A receptor (right panel).

## Results

### Retrospective docking of known ligands against the AF2 structures

We began by docking known ligands against the AF2 models of the α_2_ and 5-HT2A receptors. Our approach took two forms: (1) taking novel ligands found from docking against the experimental structures and re-docking them against the AF2 models; and (2) docking previously known ligands from the literature. Against the crystal structure of the α_2_ receptor, we had previously screened 490 million make-on-demand (“tangible”) molecules from ZINC20. From among the top-ranking 300,000 molecules (top 0.06% of the ranked library), 138 high-scoring ones were tested, of which 70 displaced over 50% of the known ligand [^3^H]-DTG at 1 µM, a hit rate of 51%^7^ (**Fig. 2a** left panel). The best 21 actives had Ki values between 1.8 nM and the low µM range, with 19 actives having Ki < 50 nM, and 6 having Ki < 5 nM^7^ (**Fig. 2b**). When we re-docked the 490 million molecule library against the AF2 model of σ_2_, the ranks of the 138 high-scoring molecules against the crystal structure dropped substantially, and were no longer among the top 470,000. This may reflect a slight shrinking of the AF2 orthosteric site versus that in the crystal structure, which militates against the binding of many of the larger ligands explored among the 138. Correspondingly, when docking known α_2_ ligands from the ChEMBL database versus a library of property matched decoys—a widely used control in docking—the crystal structure (logAUC: 39) returned much higher enrichments than did the AF2 model (logAUC: 16, **Extended Data Fig. 2a**). Similar retrospective studies against the cryoEM and AF2 structures of the 5-HT2A receptor had similar results: the experimental structure led to higher retrospective enrichment of 41 known ligands over 2050 property-matched decoys for the experimental structure than it did for the AF2 structure (**Extended Data Fig. 2b**). These observations are consistent with previous *retrospective* studies comparing AF2 models to experimental structures for ligand discovery.

### *Prospective* docking against the σ_2_ AF2 structure

A potential flaw in the retrospective logic is that the ligands chosen are biased by the receptor structure, and the receptor conformation is adapted to the ligands with which it was determined. For instance, the 70 new docking hits fit the crystal structure against which they were selected better than the AF2 structure, and even ChEMBL ligands share this “bias of the known”. Thus, it is conceivable that even though the AF2 structures are worse at recapitulating known actives, they might still may be able to prioritize new ligands from large library docking. To investigate this possibility, we docked the same 490 million molecule tangible library against the σ_2_ receptor’s AF2 model, selecting 119 molecules from the top 300,000 for synthesis and testing (**Supplementary Information Table 1**). Of these, 64 molecules displaced over 50% [^3^H]-DTG specific σ_2_ binding at 1 µM, a hit rate of 54%. While slightly higher than the 51% observed in the crystal structure docking campaign, the two hit rates were not significantly different based on a z-test. The top 18 hits from the AF2 campaign had Ki values between 1.6 nM and 84 nM, with 13 having Ki’s < 50 nM, and 2 with Ki’s < 5 nM (**Fig. 2b, Extended Data Fig. 3**). Although the highest affinity is comparable between the screens for the crystal and AF2 structures, the affinity distribution is slightly better for the crystal structure campaign. Intriguingly, despite the similarities in hit rates and affinity ranges, only four of the 125 new ligands shared the same core scaffold between the crystal structure and AF2 campaigns (**Extended Data Fig. 4**), and certainly none of the ligands between the two campaigns were the same. By ECFP4-based Tanimoto coefficient (Tc), the two sets of experimentally confirmed ligands have an average Tc of 0.32, not far from random for this fingerprint. Consistent with the diversity, the most potent ligand from the AF2 campaign, ZINC866533340 (Ki 1.6 nM), represented a chemotype previously unseen for the σ_2_ receptor (**Fig 2c** and **2d**).

**Figure 2.**
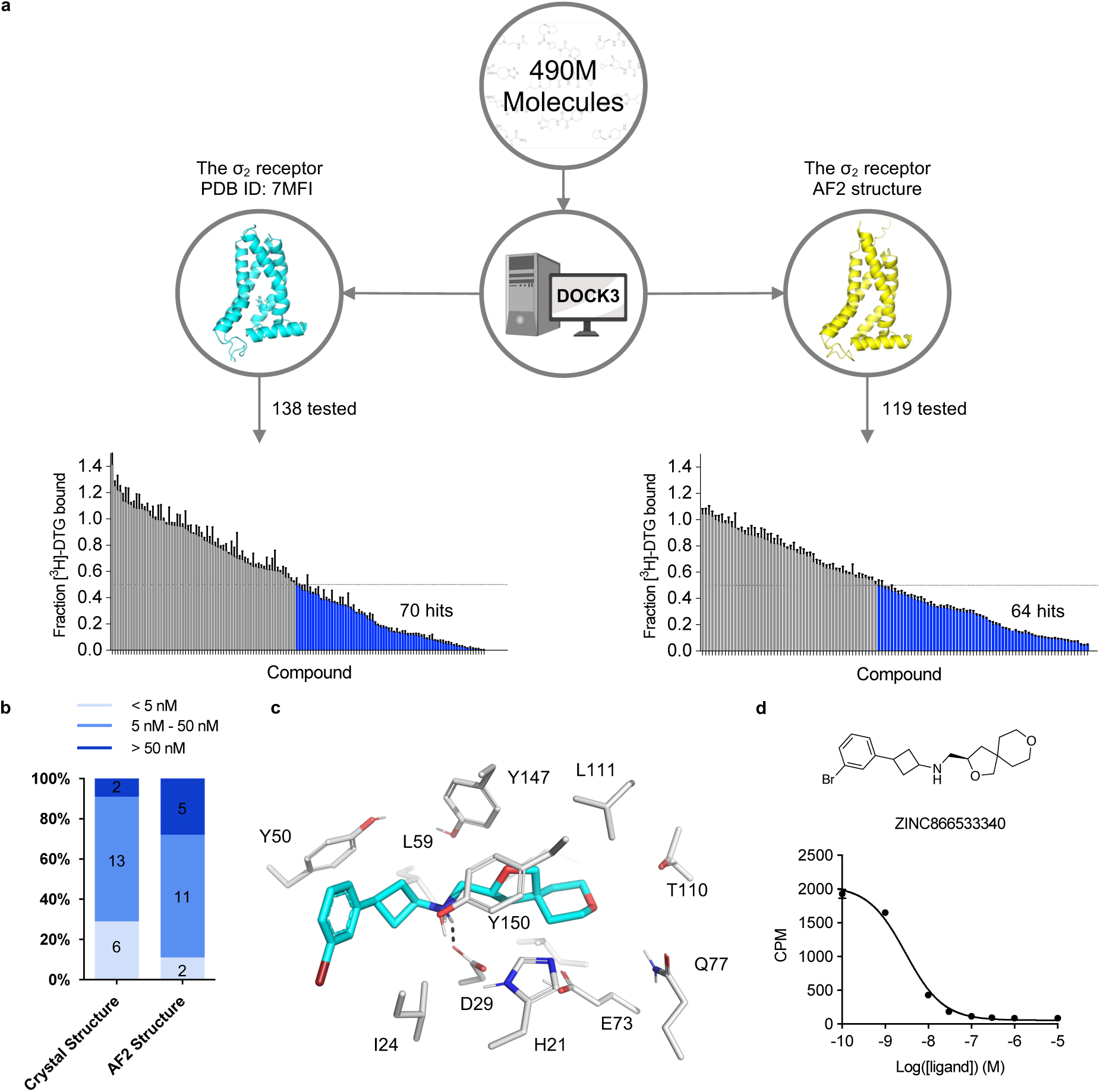
Comparison of prospective screens against the crystal and AF2 structures of the σ_2_ receptor. **a.** The same 490 million molecules from ZINC20 were screened against both the crystal and AF2 structures of the σ_2_ receptor. From these, 138 molecules from the crystal docking campaign and 119 from the AF2 docking campaign were synthesized and tested in a radioligand displacement assay. The campaign involving the crystal structure has already been published. The left panel is replotted based on the previously published data set. Displacement of the radioligand [^3^H]-DTG by each tested molecule occurs at 1 μM (mean ± s.e.m. of three technical replicates). A dashed line indicates 50% radioligand displacement. Dots below the dashed line represent confirmed binders, which are colored in blue. **b.** The distribution of binding affinity levels among the hits from both the AF2 and crystal structure screens. We measured competition binding curves for the 21 top docking hits from the crystal structure screen and the 18 top hits from the AF2 structure screen. These hits are categorized into three affinity ranges: <5 nM; 5 nM–50 nM; and >50 nM. **c.** The docked poses of the best binder from the screen against the AF2 model. **d.** The competition binding curve of the best binder from **c** against the σ_2_ receptor. The data are represented as mean ± s.e.m. from three technical replicates.

### *Prospective* screens against 5-HT2A receptor structure and AF2 model

As an experimental structure against which to dock, we used the complex of the 5-HT2A receptor with the partial agonist lisuride and with mini Gαq (**Extended Data Fig. 5**). Lisuride is a potent and non-psychedelic partial agonist, and its structure seemed suited to longstanding efforts to discover novel and non-psychedelic 5-HT2A agonists^49^. Because the structure of the Lisuride/5-HT2A/mini-Gαq structure has not been previously described, we briefly do so here. To isolate an active state heterotrimeric complex. we used a mini-GαqiN-Gβ1-Gγ2 heterotrimer co-expression system in the presence of stabilizing single-chain antibody scFv16^50^. Each component was purified separately, and the active state complex was formed in the presence of lisuride. As this approach has been widely-used to determine active-state Gαq-coupled complexes^51–54^, we then subjected this complex to single-particle cryo-EM analysis and built a consensus reconstruction at 3.1 Å through multiple steps of 3D-classification and focused refinement (**Extended Data Fig. 5).** The definition of the cryoEM density allowed us to unambiguously model lisuride within the orthosteric pocket (**Fig. 3a, Extended Data Table 1**) with the binding pose validated by Gemspot ^55^. The global structural features were consistent with previous 5-HT2A ternary complex cryoEM reconstructions and with other receptor-Gαq bound structures^51–54^. In this active-state structure, lisuride recapitulates many interactions seen in an inactive-state crystallographic complex^56^ (PDB: 7WC7), with some important differences. As in the crystal structure, lisuride’s indole hydrogen-bonds with S242^5.46^, its cationic nitrogen ion-pairs with D155^3.32^, and its diethylamide packs with W151^3.28^ (**Extended Data Fig. 6**) (superscripts use Ballesteros– Weinstein and GPCRdb^57^ nomenclature^58^). Intriguingly, lisuride’s pose within the orthosteric site in a previously reported crystal structure shifts due the presence of a penetrating lipid between TM4/5 (**Extended Data Fig. 6**), which was not present in the cryoEM structure. This lipid contributes to a rotation inward of Phe-234^5.38^ versus the cryoEM structure, which is one of the primary conformational differences in the orthosteric site between the lisuride cryoEM structure, used for the experimental structure docking, and the AF2 model (see below). We do note that the AF2 model was predicted before the lisuride-bound crystal structure was deposited in the PDB, perhaps indicating that it is capturing a low energy sidechain orientation not observed in the cryoEM reconstructions.

Unlike the case with the σ_2_ receptor, the AF2 model of the 5-HT2A receptor orthosteric site exhibited notable rotamer changes in several residues compared to the cryoEM structure. Overall, the backbone RMSD between modeled and experimental structures was 1.6 Å (compared to 0.5 Å for σ_2_). While most orthosteric residues had an RMSD < 1.5 Å, key residues Phe-234^5.38^ on TM5 and Leu-229^45.52^ on the ECL2 loop differed by 2.5 and 3.1 Å, respectively, adopting different rotamers in the two structures (**Fig.1, right panel**); Trp-151^3.28^. Moreover, Trp-151^3.28^ is pushed outward in the lisuride cryoEM structure (PDB ID: 8UWL), due to packing with the ligand’s diethylamide, relative to the AF2 model. These differences shrink the binding pocket in AF2 model versus that of the lisruide cryoEM structure used in the docking. Since the AF2 model draws on sequence similarity from an overall family of related 5-HT receptors, and because the cryoEM structure represents an active state of the receptor in complex with Gαq whereas the AF2 model more closely resembles an inactive state (**Extended Data Fig. 7**), we anticipated that the experimental structure would be more likely to select agonists that are 5-HT2A-selective and Gαq biased. On the other hand, both docking campaigns targeted residues key for agonist recognition, for bias, and for subtype-selectivity^59^ (S239, S242, T160, F234, and F339). In this sense, both docking campaigns were biased toward finding 5-HT2A agonists. Over 1.6 billion tangible molecules were docked against both the cryoEM and AF2 receptor structures using DOCK3.8. Applying consistent filtering, clustering, and hit-picking criteria, we prioritized 223 (from docking against the cryoEM structure) and 161 (from docking against the AF2 structure) top-ranking and internally diverse molecules — all within the top 3 million (top 0.2%) of docking-ranked molecules — for synthesis and testing (**Supplementary Information Table 2**).

In primary radioligand binding screening assays, 52 of the 223 molecules from the cryoEM docking displaced over 50% [^3^H]-LSD at 10 µM of the docked ligand (**Fig. 3a** and **3b**, left panel), a hit rate of 23%. Meanwhile, 42 of the 161 molecules from the AF2 docking campaign met this threshold, a hit rate of 26% (**Fig. 3a** and **3b**, right panel). Applying a stricter criterion, where displacement over 90% [^3^H]-LSD at 10 µM defines a potentially potent hit, the rates were 4% (8 hits/223 tested) and 6% (9 hits/161 tested) for the cryoEM and AF2 docking, respectively. In secondary radioligand binding assays, these top 17 hits had Ki values between 15 nM and 344 nM (**Extended Data Fig. 8** and **Extended Data Table 2).** Unexpectedly, the three compounds with the highest affinity (15 to 24 nM) were all from the AF2 docking campaign, while the three best compounds from the cryoEM docking campaign displayed affinities about five-fold weaker (between 71 to 114 nM) (**Extended Data Fig. 8 and Extended Data Table 2**). While the AF2 docking hit rates are higher, they do not differ significantly from the experimental docking hit rates based on a z-test. Despite the similar hit-rates, the chemical identities of the active molecules differed. Of the 95 active molecules from both campaigns, no two shared the same scaffold (**Extended Data Fig. 4,** bottom panel), and the Tc average pairwise similarity of 0.27 close to random.

**Figure 3.**
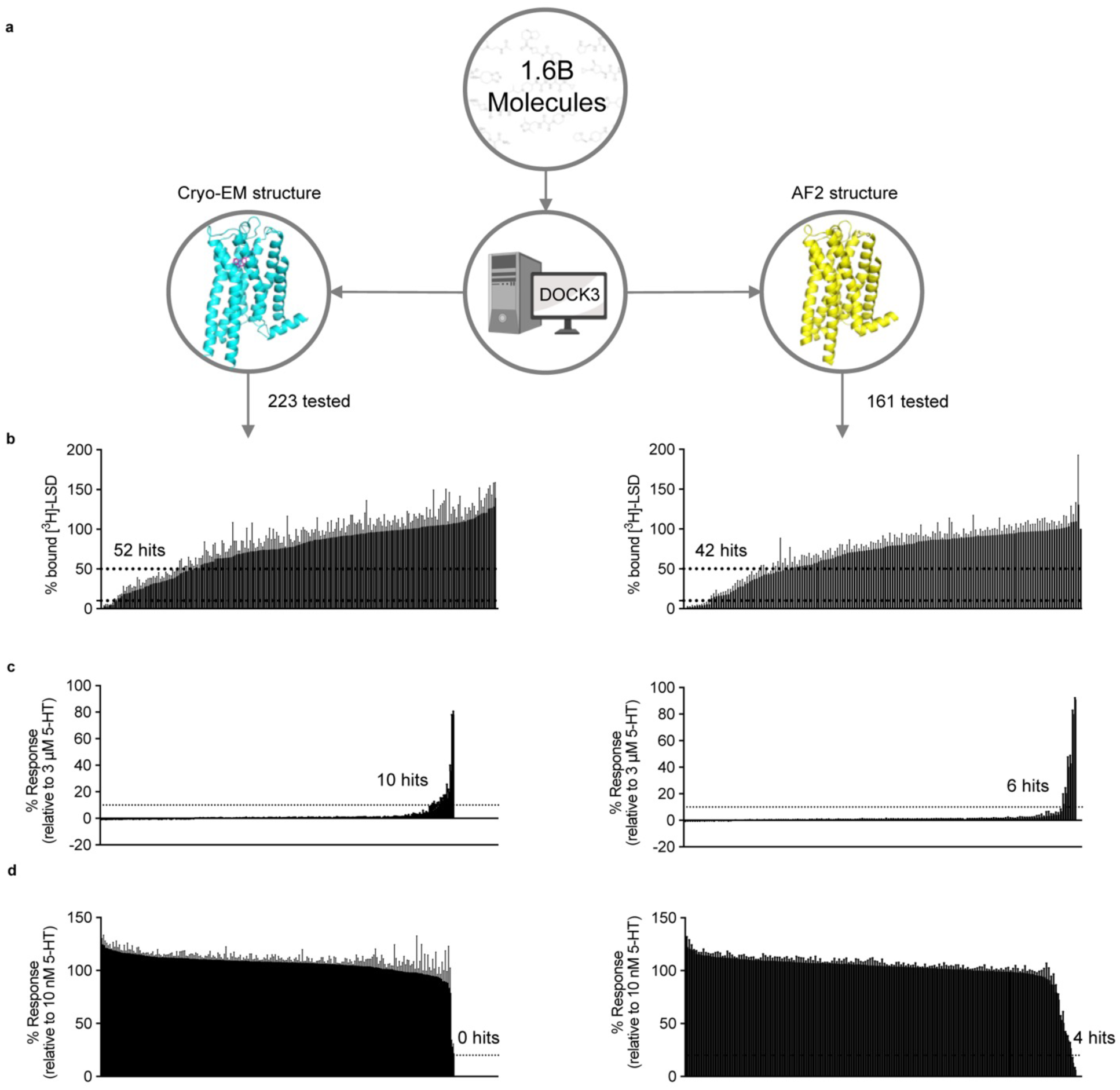
Comparison of prospective screens against the cryoEM and AF2 structures of the 5-HT2A receptor. **a**. The same set of 1.6 billion molecules from ZINC22 were docked against the cryoEM and AF2 structures of the 5-HT2A receptor. 223 molecules were prioritized from the cryoEM docking campaign (left) and 161 from the AF2 docking campaign (right). **b.** Displacement of the radioligand [^3^H]-LSD by each molecule at 10 μM (mean ± s.e.m. of three independent replicates). Dashed lines indicate 50% and 90% radioligand displacement respectively. **c.** The Ca^2+^ mobilization functional assay in agonist mode. Each compound was tested at a concentration of 3 μM. A dashed line indicates agonism equivalent to 10% 5-HT activity. Data are presented as mean ± s.e.m. from three biological replicates. **d.** The Ca^2+^ mobilization functional assay in antagonist mode. Each compound was tested at a concentration of 3 μM. A dashed line indicates antagonism equivalent to 20% clozapine activity. Data are presented as mean ± s.e.m. from three biological replicates.

We next screened the 338 docking-prioritized molecules from both campaigns for functional activity, for sub-type selectivity among the 5-HT2A, 5-HT2B, and 5-HT2C receptors, and for ligand bias between Gαq activation or β-arrestin2 recruitment. Following treatment with 3 μM of each compound, 10 compounds from the cryoEM docking set and 6 from the AF2 docking set emerged as 5-HT2A agonists, defined by ≥ 10% 5-HT response (**Fig. 3c**). We also observed 10 compounds (5 cryoEM and 5 AF2) and 6 compounds (3 cryoEM and 3 AF2) that were 5-HT2B and 5-HT2C agonists (**Extended Data Fig. 9**). When screened for antagonist activity at 3 μM, 4 compounds (0 cryoEM and 4 AF2) antagonized 5-HT2A receptor activity, as defined by ≥ 20% clozapine activity (**Fig. 3d**). Additionally, we found 17 compounds (10 cryoEM and 7 AF2) and one compound (cryoEM) with antagonist activity at the 5-HT2B and 5-HT2C receptors, respectively (**Extended Data Fig. 9**). Interestingly, comparisons between our primary binding (**Fig. 3b**) and functional screening data (**Figs. 3c** and **3d**) indicates that two of the 17 top binding hits exhibit 5-HT2A agonist activity (**Extended Data Fig. 8 and Extended Data Table 2**), while 15 antagonized 5-HT2A activity. Remarkably, both of the high affinity agonists (Z2504 and Q2118) were from the AF2 dataset. Thus, more agonists were found from docking to the cryoEM structure than to the AF2 model, and antagonists were only found from docking against the AF2 structure. Thus far, this is consistent with docking against the activated conformation represented by the cryoEM structure and docking against an inactive conformation represented by the AF2 structure (**Extended Data Fig. 7**).

Dose-response curves measuring calcium mobilization confirmed 5-HT2A agonist activity of the top hits from both the cryoEM and AF2 docking sets (**Fig. 4a** and **4b**, top panels). The potencies (pEC_50_ values) of top agonists from the cryoEM docking set ranged from 246 nM to 3 μM while those from the AF2 docking set ranged from 42 nM to 1.6 μM (**Fig. 4** and **Extended Data Table 3**). Three of the top five AF2 agonists (Q2118, Z7757, and Z2504) displayed subtype-selectivity for 5-HT2A over the 5-HT2B and 5-HT2C receptors (**Fig. 4b**). Meanwhile, none of the top cryoEM agonists displayed any degree of 5-HT2A receptor selectivity.

In BRET experiments measuring heterotrimeric Gαq protein dissociation, which reflects G protein activation,^60^ versus β-arrestin2 recruitment (**Extended Data Fig. 10, Extended Data Table 3**), nine of the top 10 agonists from both docking sets exhibited modest bias toward Gαq signaling versus 5-HT, while only one compound showed modest bias toward β-arrestin2 recruitment at high concentrations. Similar experiments with the top antagonist hits identified from primary (**Extended Data Fig. 9**) and functional screening (**Extended Data Fig. 11**) confirmed weak inhibition of 5-HT2A receptor activity, with potency of antagonists from the cryoEM docking set between 13 μM and 78 μM, while those from the AF2 docking set were between 907 nM and 114 μM (**Extended Data Table. 4**).

**Figure 4.**
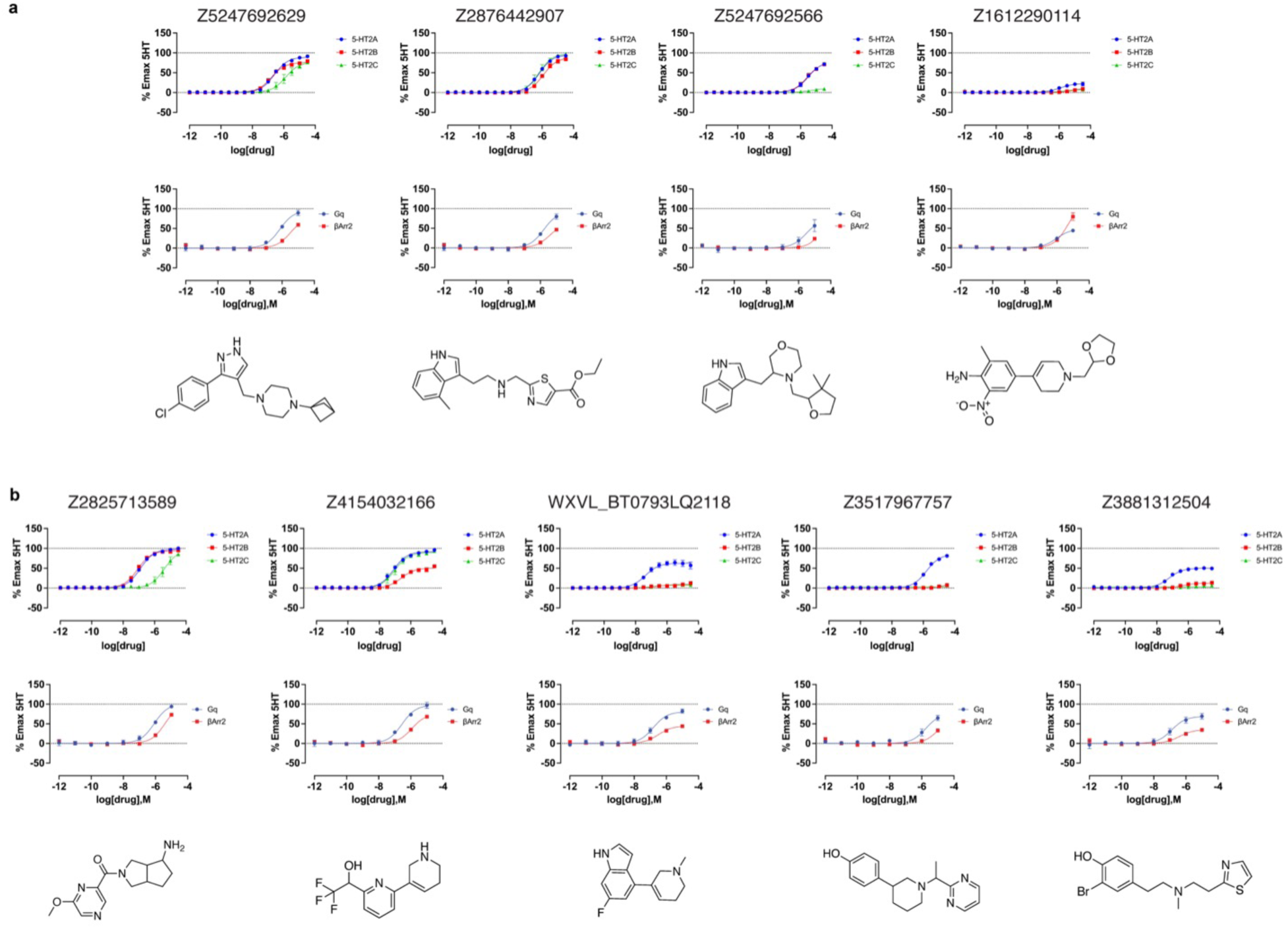
Dose-response curves of the top agonists against 5-HT2A, 5-HT2B, and 5-HT2C. Top agonists from both docking campaigns were tested at the 5-HT2A (blue), 5-HT2B (red), and 5-HT2C (green) receptors. **a.** Functional assays measuring calcium mobilization (top) and Gαq protein dissociation or β-arrestin2 recruitment (bottom) for the top4 agonists from the cryoEM docking campaign. **b.** Functional assays measuring calcium mobilization (top) and Gαq protein dissociation or β-arrestin2 recruitment (bottom) for the top5 agonists from the AF2 docking campaign. The chemical structure of each compound is displayed below its respective dose-response curve. Data are presented as mean ± s.e.m. of three biological replicates.

We next determined a cryoEM structure with one of the 5-HT2A selective agonist from the AF2 screen, Z7757 in complex with 5-HT2A and mini-Gαq (**Fig. 5b**), which by chemotype was the most unusual of the new agonists, with little topological similarity to 5-HT2A receptor agonists of which we are aware. Here the 5-HT2A receptor was co-expressed with mini-GαqiN-Gβ1-Gγ2 heterotrimer and purified in the presence of Z7757 and scFv16, as previously described ^53,61,62^. This permitted direct isolation of the complex and after a single size-exclusion step allowed us to use a large amount of complex in grid preparation. Purified complexes were subjected to single-particle cryoEM analysis (see **Extended Data Fig. 13** for processing tree), where particles that did not contain scFv16 were filtered out by 2D/3D classification and a focused no alignment 3D-classification and refinement on the receptor was carried out). The structure was ultimately determined to a global resolution of 3.0Å, allowing us to unambiguously build Z7757 within the orthosteric pocket (**Fig. 5b**). The binding pose was further validated utilizing Emerald (**Extended Data Fig. 14**), which uses a genetic algorithm to fit the ligand using the cryoEM density map as restraints^63^. The Cα-RMSD of the receptor was close to the lisuride structure (0.78Å) and recapitulated the receptor-Gαq interactions in prior 5-HT2AR structures (**Fig. 5c**) ^49,53,61,64^.

The binding of Z7757 in the 5TH2A cryoEM structure closely resembles the predicted docking pose. As anticipated in the docked model, Z7757 interacts with key recognition and activation residues (**Fig. 5d-f**); the docking prediction of Z7757 superposes on the experimental result with an RMSD of 1.6Å with the major difference between the two coming from a 44° rotation of the agonist’s distal pyrimidine ring (**Fig. 5g**). In both the docking prediction and the cryoEM structure, the new agonist uses its phenolic hydroxyl to hydrogen bond with well-known activation residues on TM3 and TM5 (T160^3.37^ and S242^5.46^), and also with the backbone of G238^5.42^ an interaction predicted in the docking but unusual among 5-HT2A receptor agonists. As expected, the cationic nitrogen of Z7757 ion-pairs with the key recognition D155^3.32^ of 5-HT2A, and packs well with the binding site in both the modeled and the experimental structures.

Stepping back from the local interactions between Z7757 and receptor, it was interesting to consider how the model compares to the experimental result in regions where we expected important differences: the positions of Leu-229^45.52^, Phe-234^5.38^, and Trp-151^3.28^ which adopt different conformations and even rotamers in the AF2 model and in the lisuride/5-HT2A receptor structure. The rotamer of Phe-234^5.38^ in the Z7757-bound structure closely resembles that in the lisuride-bound structure (**Fig. 5i**). While this is an important difference which the AF2 model failed to anticipate, in the Z7757 complex F234^5.38^ is relatively distant from the new agonist, with their closest approach being 5.9Å away, reducing the impact of this residues on recognition. Intriguingly, the overall position of Leu-229^45.52^ in the Z7757 complex more closely resembles that of the AF2 model than that of the experimental structure used for docking (**Fig. 5i**). Finally, W151^3.28^ in the Z7757 complex more closely resembles the lisuride cryoEM structure used in the experimental structure docking than the AF2 model. Nevertheless, W151^3.28^ has still swung inward relative to its position in the lisuride cryoEM complex used for docking—in the AF2 model this pocket-constricting rotation is further exaggerated. The constriction of this part of the orthosteric site reduces the complementarity of LSD-like ligands, including lisuride and many others characterized for the 5HT2A receptor, while still allowing the pyrimidine of Z7757—and perhaps other agonists structurally unrelated to classic 5HT2A receptor agonists—to still fit well (**Fig. 5i**). Indeed, among the 5-HT2A structures deposited in the PDB, W151^3.28^ adopts a range of conformations, opening and closing this part of the site, with the active-state lisuride complex adopting the most open position of the set (Extended Data Fig. 15a-c). Meanwhile, both a non-classical agonist (*R*-69, PDB: 7RAN) (Extended Data Fig. 15b) and an antagonist (Zotepine PDB: 6A94) exhibited rotamers like the AF-model (Extended Data Fig. 15c). We also found that all the crystal structure ligand complexes with 5HT2A receptor, except that with LSD (PDB: 6WGT), have Phe-234^5.38^ in a conformation akin to that of the AF2 models used for the docking, while the cryoEM structures have consistently exhibited the altered orientation (Extended Data Fig. 15a). Taken together, these observations support the idea that the AF2 model has sampled low energy states of these residues, even though they differ from the particular experimental structure used in the docking campaign (the cryoEM structure with lisuride).

**Figure 5.**
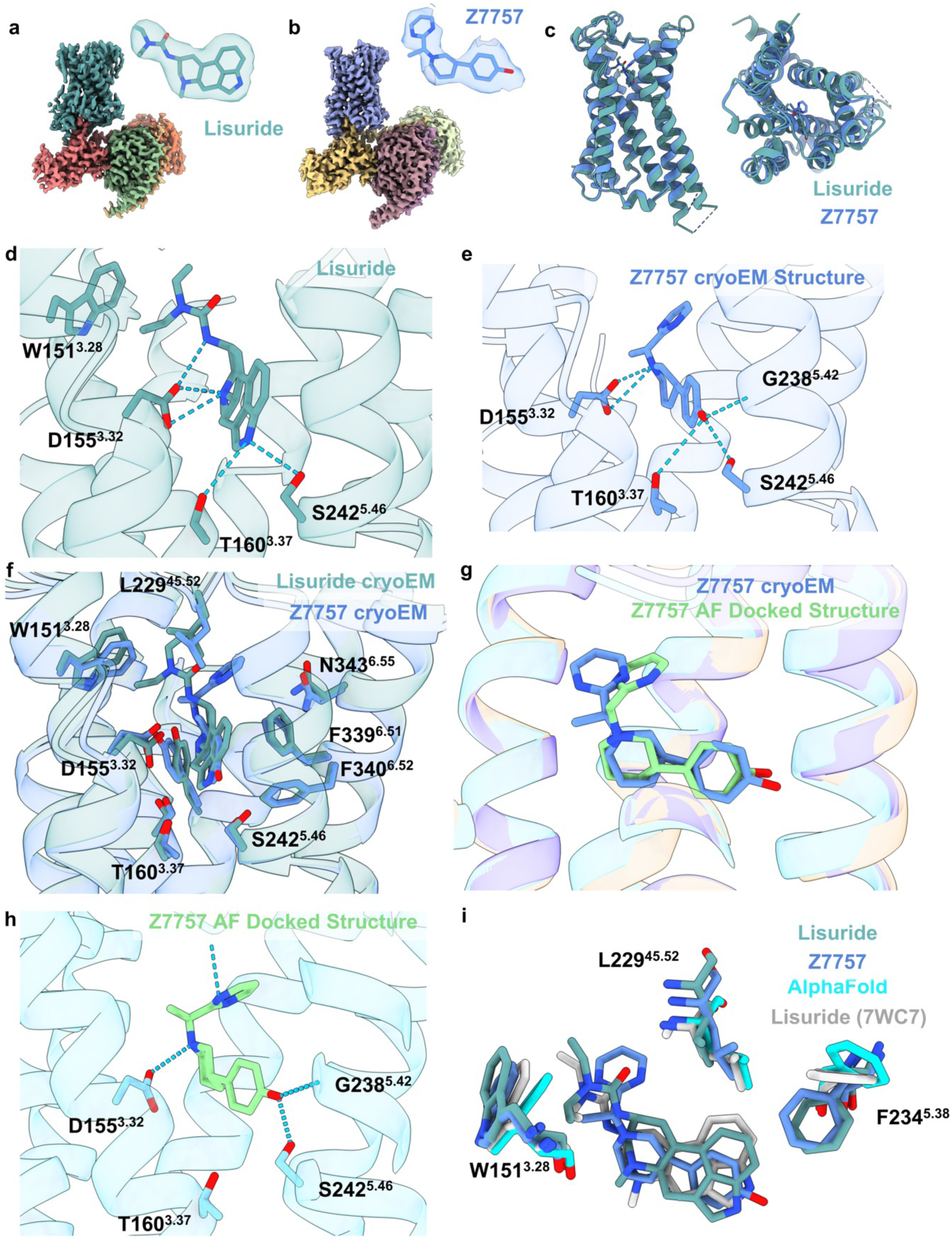
Structural Characterization of the 5-HT2A receptor in complex with Lisuride and Z7757. Maps of Lisuride (**a**) and Z7757 (**b**) active state heterotrimers respectively. Models of the compounds built into the electron density are shown. (**c**) Overlay of the Lisuride and the Z7757 structures, which superpose to 0.78 Å Cα-RMSD. (**d**) Interactions between Lisuride and (**e**) between Z7757 and the receptor. (**f**) Overlay of Z7757 and Lisuride in the orthosteric pocket. **(g)** Comparison of the experimental Z7757 structure and the predicted structure from the AF2 docking screen. (**h**) Predicted interactions from the docked pose of Z7757 in the AF2 structure. (**i**) Aligned Lisuride cryoEM structure, AF2-model, Z7757 cryoEM structure, and the Lisuride crystal structure (7WC7) highlighting residues that showed the biggest difference between the AF2-model and Lisuride cryoEM structures and that were used in the docking studies.

## Discussion

Our main finding is that, unexpectedly, the prospective large library docking campaigns against the AF2 models were no less effective than those against experimental structures. The hit rates were high for both the 02 and the 5HT2A receptors across hundreds of molecules experimentally tested against each model for both targets, and were not significantly different between the modeled and experimental structures. For the σ_2_ receptor, 54% of the AF2-derived docking hits were active at 1 µM, while for the crystal structure-derived docking the hit rate was 51%—not statistically different (**Figure 2a**). Meanwhile, for the 5-HT2A receptor, 26% of the molecules from the AF2-derived model bound at 10 µM, while for the cryoEM experimental structure 23% bound (**Figure 3b**). A comparison of the affinities of these molecules yielded similar conclusions. Against the 0_2_ receptor, the top 18 hits from the AF2 docking had K_i_ values between 1.6 nM to 84 nM. While the distribution of these affinities was detectably worse than those docked against the crystal structure (**Figure 2b**), the differences were small. Against the 5-HT2A receptor, the AF2 structure led, if anything, to more potent and selective compounds. The most potent AF2-derived agonists had EC_50_ values ranging between 42 nM and 1.6 µM, while for the cryoEM-derived docking hits the EC_50_ values ranged from 246 nM to 3.0 µM (**Figure 4**). Whereas three of these five AF2-derived docking hits had substantial sub-type selectivity (>6 to 278-fold selectivity for 5-HT2A over 5-HT2B and 5-HT2C), the cryoEM-derived docking hits were not sub-type selective. Indeed, the AF2-derived docking hits are among the most potent and selective molecules to emerge from extensive structure-based campaigns^49^ against this target.

A cryoEM structure of one of the agonists from the AF2 model docking, Z7757, largely confirmed the modeled prediction: the docked ligand superposed with its experimental pose with an RMSD of 1.6 Å (**Fig. 5g**), and the experimental structure recapitulated the hydrogen bonds anticipated in the docking. Even some of key structural features of the AF2 model were retained in the Z7757 experimental complex. For instance, of the three residues with substantial rotameric differences between the AF2 model and the lisuride structure used for docking, the position of Leu-229^45.52^ ultimately resembled that of the AF2 model in the Z7757 complex. These findings are consistent with the idea that the AF2 model captured low energy, accessible states of the 5-HT2A receptor, even though those were not seen in the experimental structure used for docking. Overall, for targets where the AF2 models resemble to the experimental orthosteric sites, such as the σ_2_ receptor, or similar but with some meaningful differences (5-HT2A receptor), structure-based screens against AF2 models can be productive.

These prospective results may be reconciled with previous retrospective studies that have suggested that experimental structures are superior to unrefined AF2 models for docking^7,39–48^. In the previous work, known ligands were docked against experimental and AF2 structures, and the former much better enriched the knowns than did the AF2 models. This is also what we observed when we docked known ligands against the σ_2_ and 5-HT2A receptors (**Extended Data Fig. 2**). However, retrospective campaigns are biased by past results, and cannot be expected to necessarily predict prospective results. Experimental structures are often determined with some of these known ligands, and certainly it may be that ligands discovered against one conformation of a protein—as in our previous campaigns against the experimental structures—might not rank as well against another low energy conformation of that same protein. The prospective results reported here, where hundreds of topologically novel chemotypes were synthesized and experimentally tested in all four campaigns (two against the experimental structures, two against the AF2 structures), support this idea. It is also supported by the low overlap, even at a scaffold level, between the active molecules found by the AF2 and the experimental docking campaigns (**Figs. 2, 3 and Extended Data Fig. 4**). The conformations of the AF2 models are, from a ligand discovery standpoint, meaningfully different from the experimental structures, and prioritize not only different molecules but different families of molecules. At least for a subset of targets, the different conformations sampled remain relevant to ligand discovery and may complement experimental structures for such purposes.

Certain caveats merit mentioning. We have focused on two targets where the AF2 conformations of the ligand binding sites are either quite close to that of the experimental structure, as for the σ_2_ receptor, where all residues adopted similar rotamers in both the AF2 model and the experimental structures, or close with a few important divergences, as for the 5-HT2A receptor, where L229 and F234 adopted different rotamers between the model and the experimental structure. There are other AF2 models that differ so much from the experimental structures that we deem them poor candidates for docking (**Extended Data Figure 1**). We have not explored how many proteins fall into these different categories, nor is it entirely clear how one would know this *a priori*. The average pLDDT scores, a measure of local accuracy in AF2 models, at the ligand binding site of MRGPRX4, which we deemed unsuitable for docking, for the 5-HT2A and for the σ_2_ receptors is 87, 95 and 95, respectively, suggesting that pLDDT scores might help rule out AF2 structures for large library docking, but this remains an active area of research. We note that the hit rates in 5-HT2A receptor functional assays are low, in the 1-5% range. Since the hit rate for binding was much higher (23 to 26%), we suspect that the low functional hit-rates may reflect, at least in part, our testing of enantiomeric mixtures of the compounds; we have found that testing pure enantiomers is often crucial for functional activity.^49^ Finally, The AF2 models used here were predicted without any ligand information. Very recent tools like RoseTTAFold All-Atom^65^ and the AlphaFold Latest^66^ can co-fold proteins with small molecules, potentially providing improved models for large-library docking.

These caveats should not obscure the central observations from this study. While *retrospective* docking of known ligands against the AF2 models of the σ_2_ and 5-HT2A receptors returned much worse enrichment than did the experimental structures (**Extended Data Fig. 2**), when large libraries were docked *prospectively* against unrefined AF2 models the hit rates were comparable to that returned by the experimental structures, and in both cases were high. Intriguingly, for both targets the AF2 models prioritized different chemotypes than did the experimental structures. This suggests that the alternate conformations sampled by AF2 models capture low-energy conformations, useful for identifying novel ligands. In the docking campaign against the 5-HT2A receptor, the functional profiles of hits derived from AF2 docking were no worse than those from experimental structure docking. Notably, three 5-HT2A receptor hits from the-AF2 docking were subtype selective, something not seen in the campaign against the cryoEM structure. Lastly, a cryoEM structure for one of the AF2-derived agonists supported the docking prediction and, indeed, certain details of the AF2 modeled receptor conformation. These results suggest that, in the right circumstances, AF2 models can much expand the range of proteins targeted for structure-based discovery, and the breadth of potent chemotypes discovered.

## Extended data figures and tables

**Extended Data Fig. 1.**
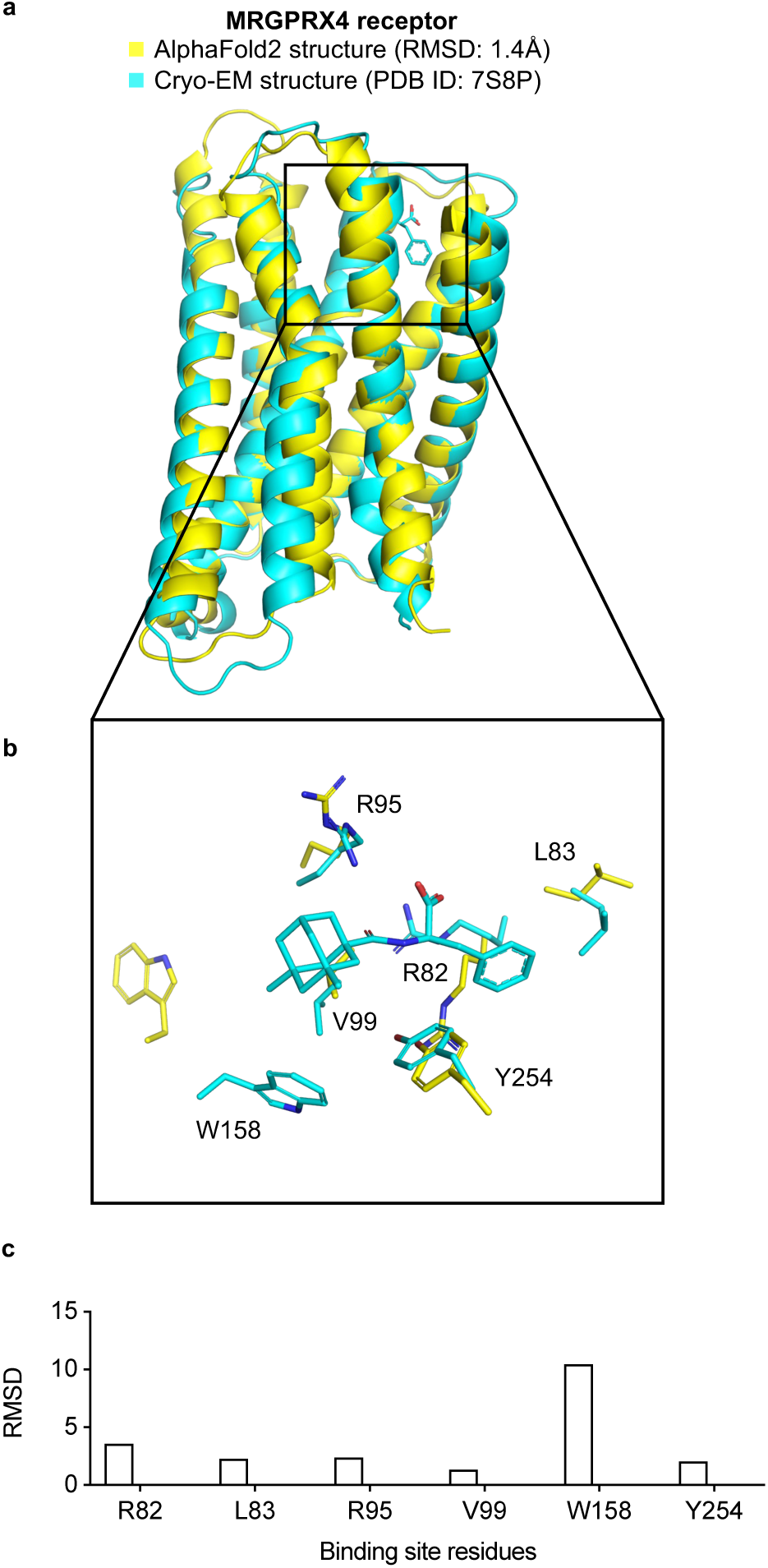
The MRGPRX4 receptor AF2 model is likely not a good candidate for docking. **a.** The experimental structure (in cyan) is overlaid with the AF2 predicted structure (in yellow). The Root Mean Square Deviation (RMSD) value is calculated based on backbone atoms. The ligand binding site residues were selected within 4 Å distance from the ligand. **b.** The full-atom RMSD values of the binding site residues between the AF2 and the experimental structures.

**Extended Data Fig. 2.**
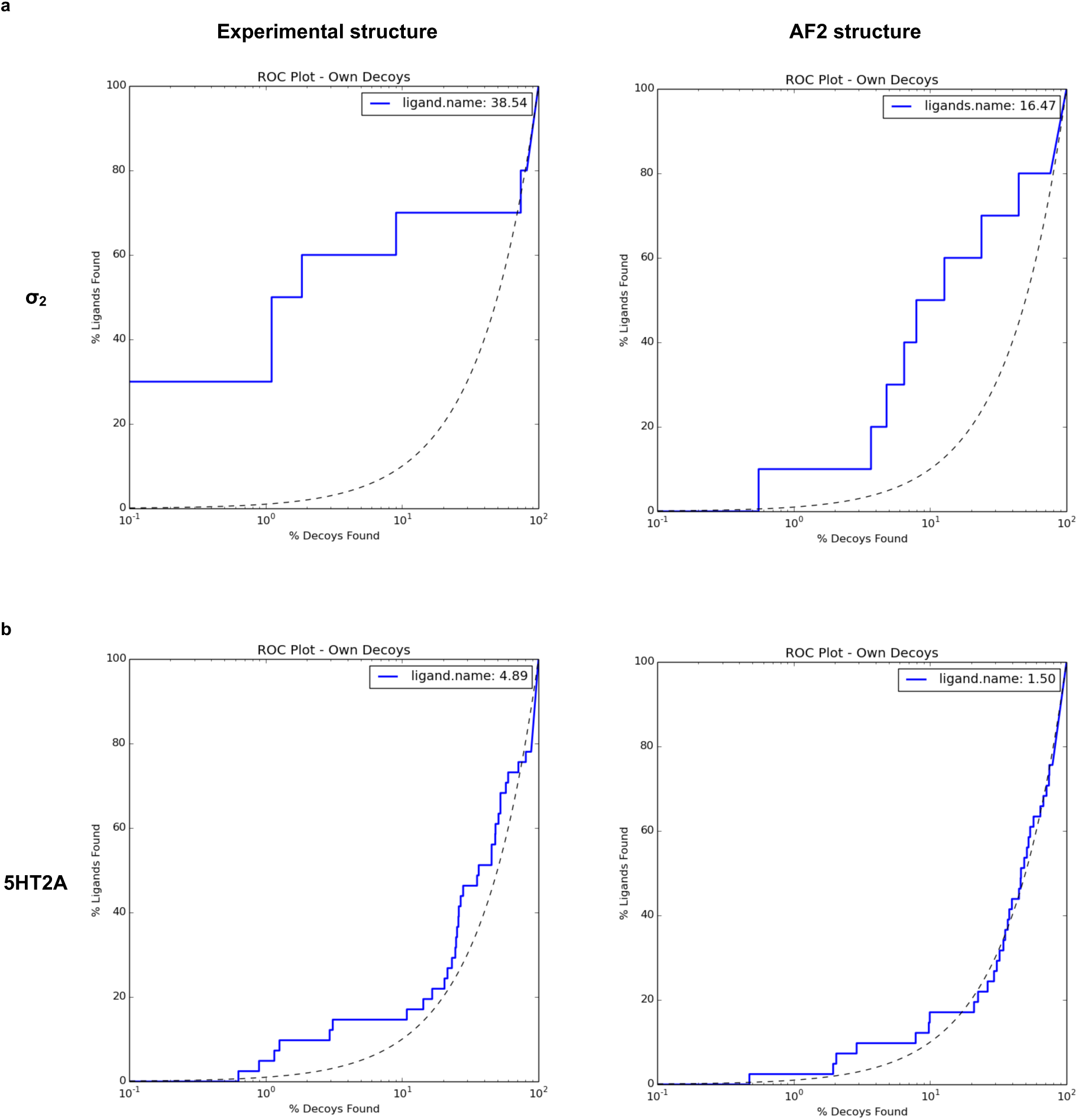
Retrospective docking of known ligands against experimental structures (left column) outperforms that for AF2 structures (right column). Panels **a** and **b** show log-transformed ROC plots that compare the rate of ligand identification versus property-matched decoys for the σ_2_ receptor and the 5-HT2A receptor, respectively. A random selection corresponds to the dashed black line. The area under this dashed line is subtracted from the reported LogAUC values. As a result, a curve above the line indicates a positive LogAUC, a curve below the line signifies a negative LogAUC, and a curve following the dashed line represents a LogAUC value of zero. In both instances, the overall LogAUC value is better when docking against experimental structures.

**Extended Data Fig. 3.**
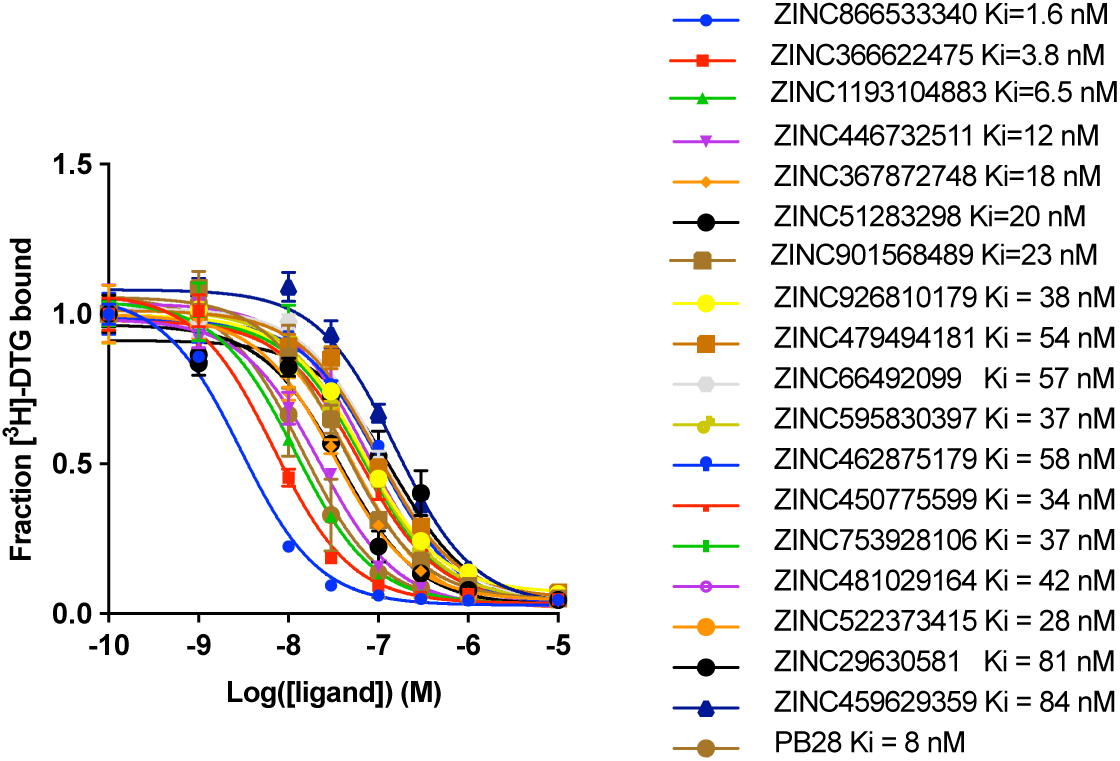
Competition binding curves for the top 18 hits from AF2 docking against the σ_2_ receptor. The data represent the mean ± s.e.m. from three technical replicates.

**Extended Data Fig. 4.**
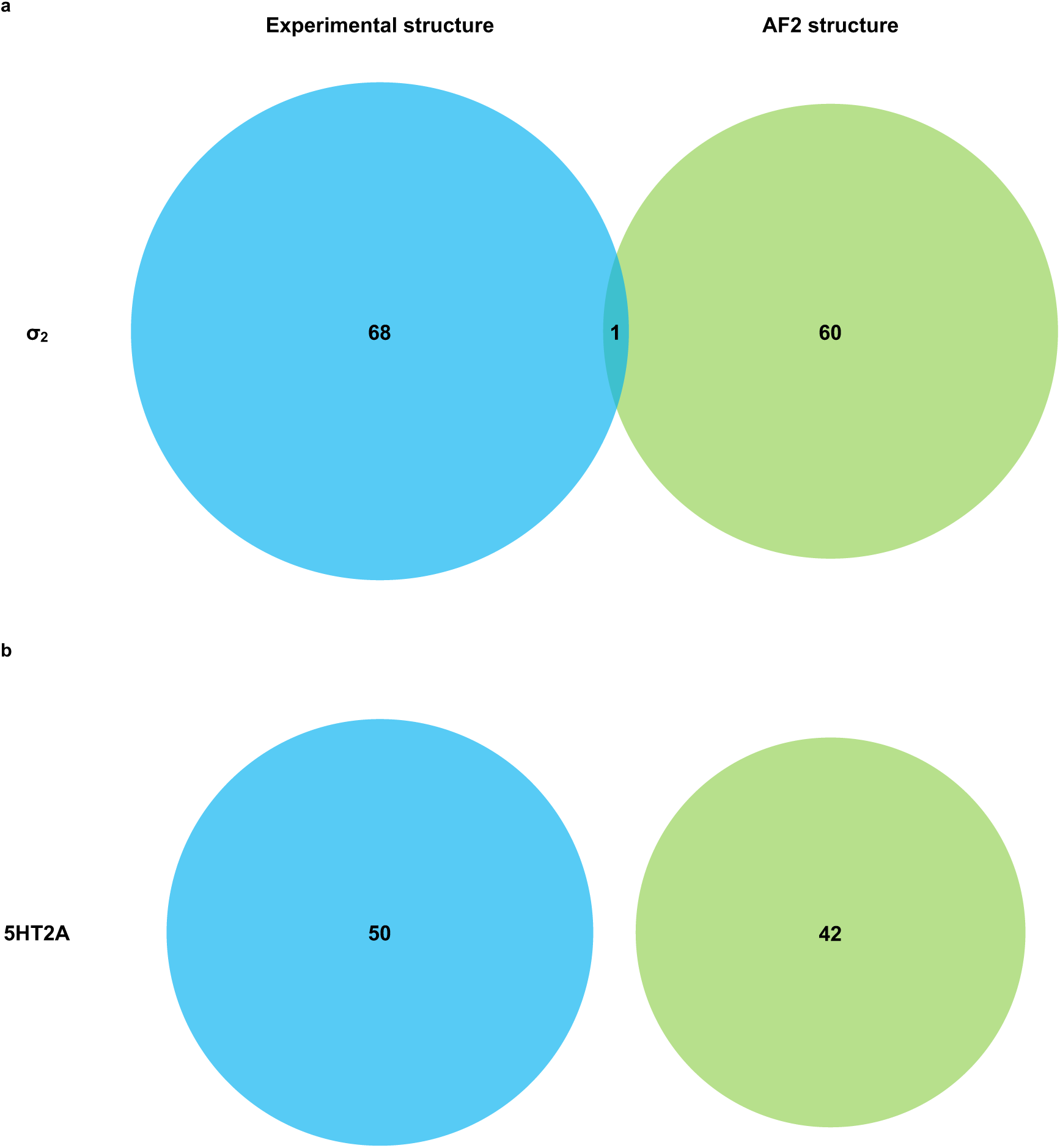
Overlap of Bemis-Murcko scaffolds between hits from docking against experimental structures (blue) and hits from docking against AF2 structures (green). Panels **a** and **b** display Venn diagrams showing scaffold overlap for hits from the σ_2_ receptor and the 5-HT2A receptor, respectively.

**Extended Data Fig. 5.**
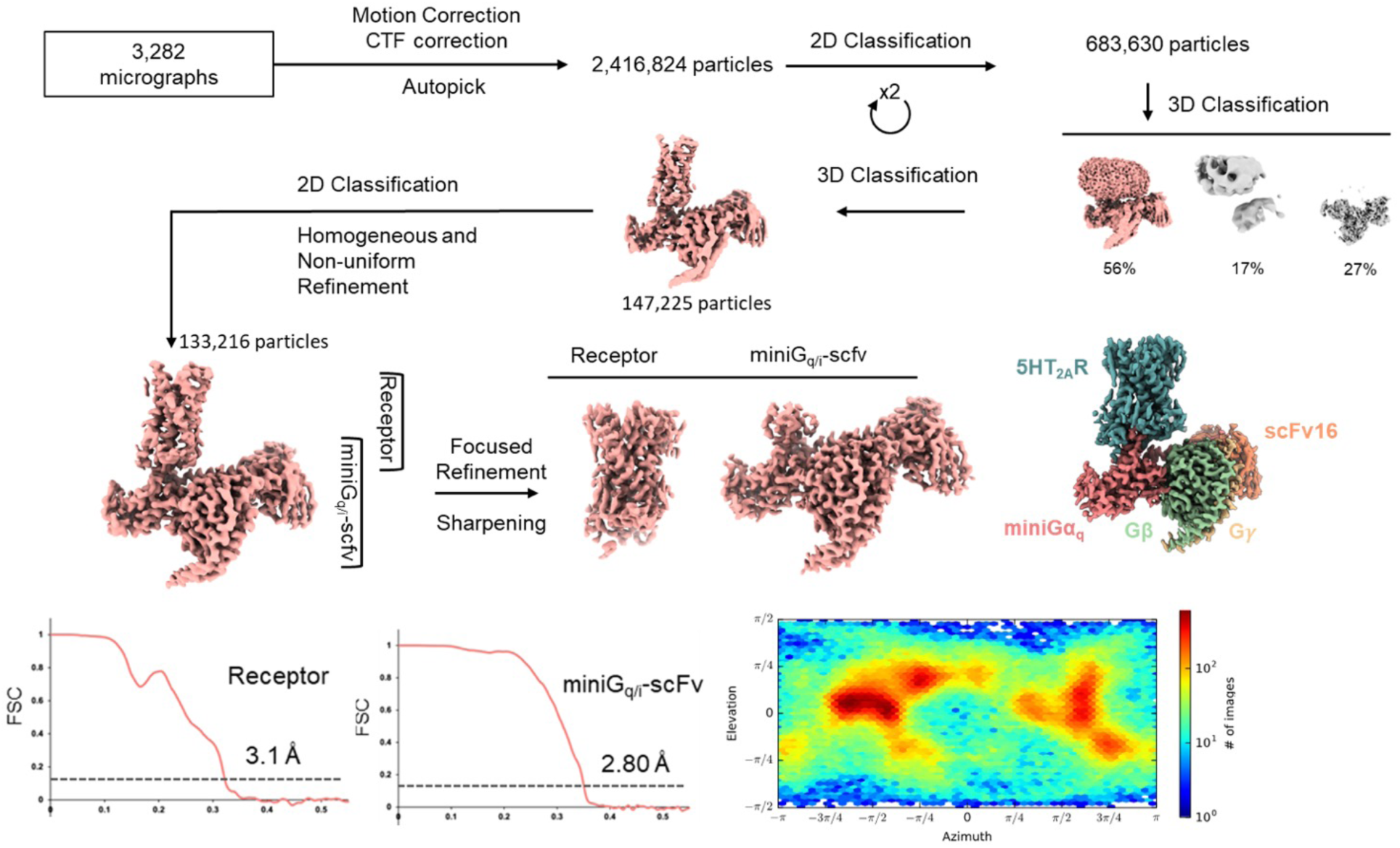
cryoEM processing Flow Lisuride. Comprehensive processing flow for the lisuride structure. After 2D-classification and several rounds of 3D classification, a focused refinement was done on the receptor and mini-Gq heterotrimer/scFv16 separately. These were then combined in Chimera. FSC plots for the receptor and heterotrimer as well as an angular distribution plot are also shown.

**Extended Data Fig. 6.**
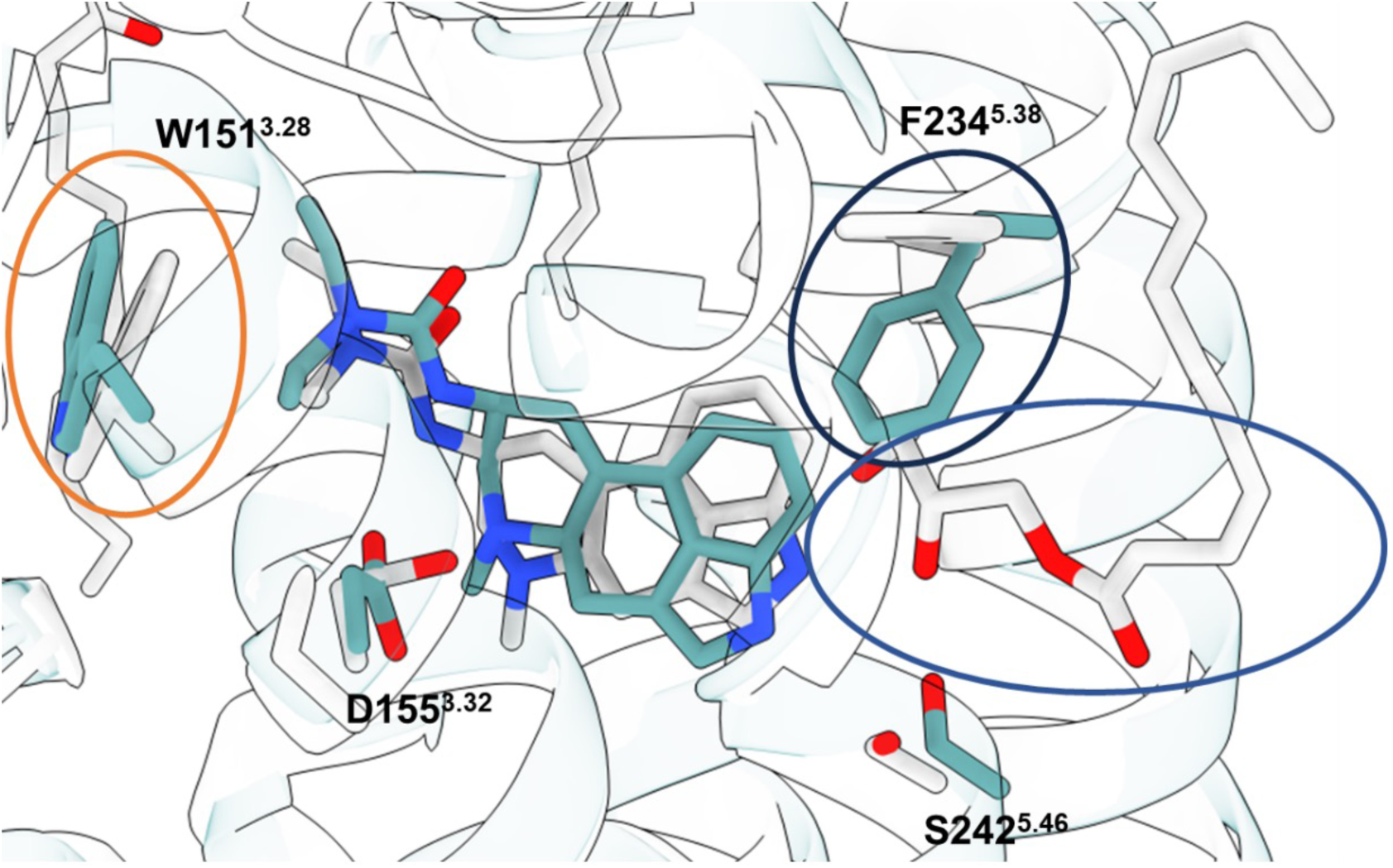
Comparison of 5-HT2A lisuride cryoEM structure and crystal structure. Overlay of the lisuride cryoEM structure solved in this work and the previously published 5-HT2A-lisuride crystal structure (PDB ID: 7WC7). There are some differences within the orthosteric pocket, but small shifts are noticed between the two structures. W151^3.28^ (orange circle) shows a small shift inward, but also follows the slight shift of lisuride within the binding pocket. Additionally due to the penetration of a lipid moiety (blue circle) into the orthosteric pocket, F234^5.38^ also was changed significantly (black circle). The pose found in the crystal structure more closely resembles the predicted structure from AF, indicating it may be sampling this space.

**Extended Data Fig. 7.**
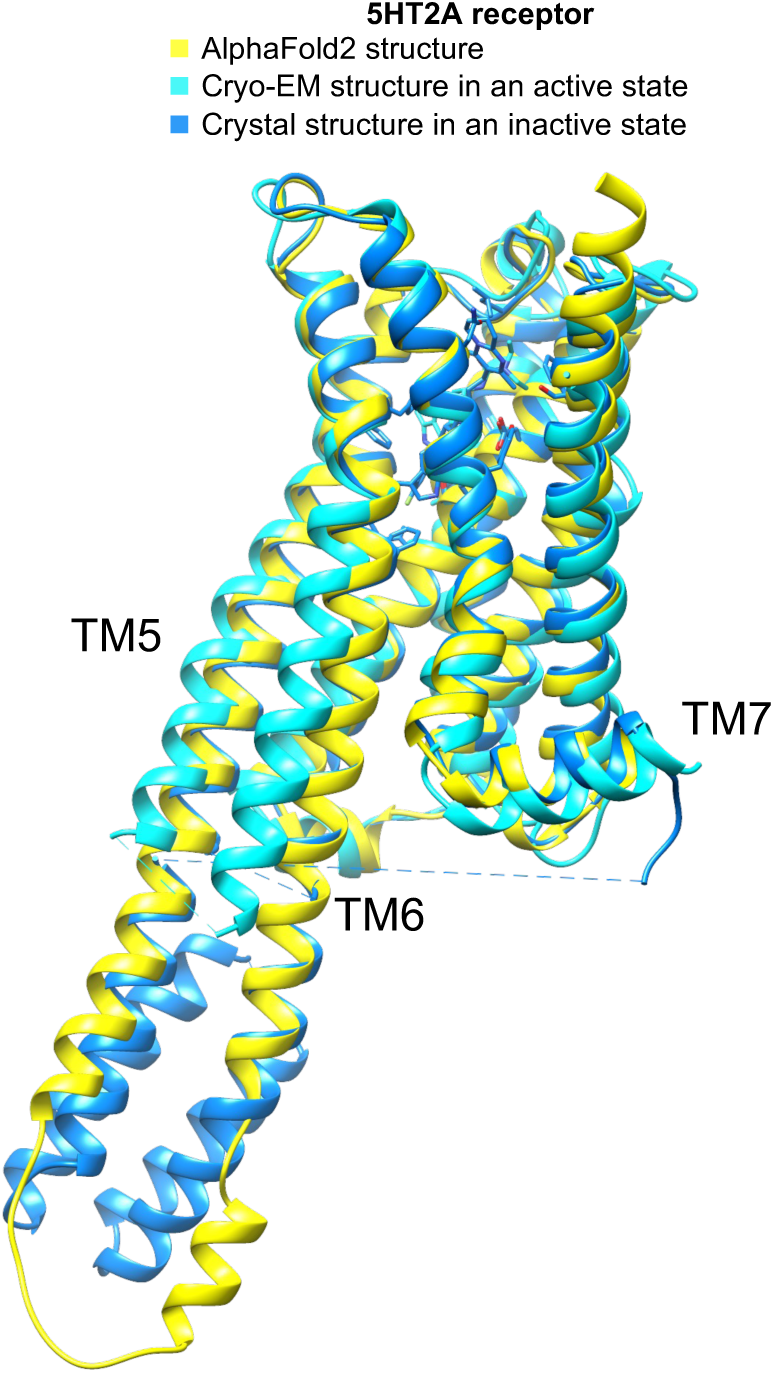
The overall AF2 structure of the 5-HT2A receptor used in this study leans more towards an inactive state. The AF2, Cryo-EM structure in an active state (PDB ID: unpublished), and the crystal structure in an inactive state (PDB ID: 6A93) are depicted in a cartoon representation and are colored in yellow, cyan, and marine, respectively. The transmembrane helix TM6 from the AF2 structure aligns more closely with that of the crystal structure in its inactive state than with the Cryo-EM structure in its active state.

**Extended Data Fig. 8.**
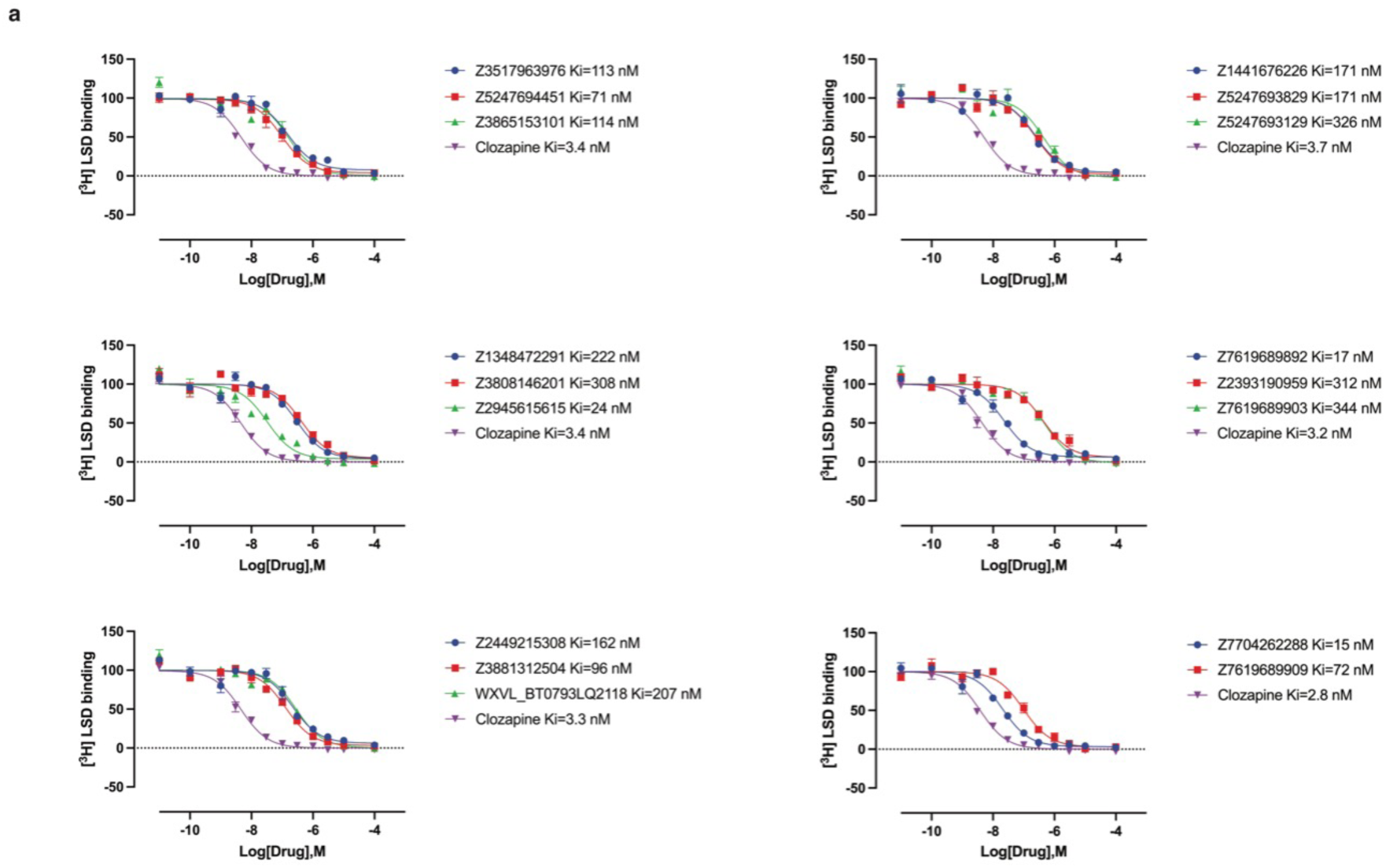
Competition binding curves for ligands that displace ≥ 90% [3H]-LSD at the 5-HT2A receptor. These data represent the mean ± s.e.m. from three independent replicates.

**Extended Data Fig. 9.**
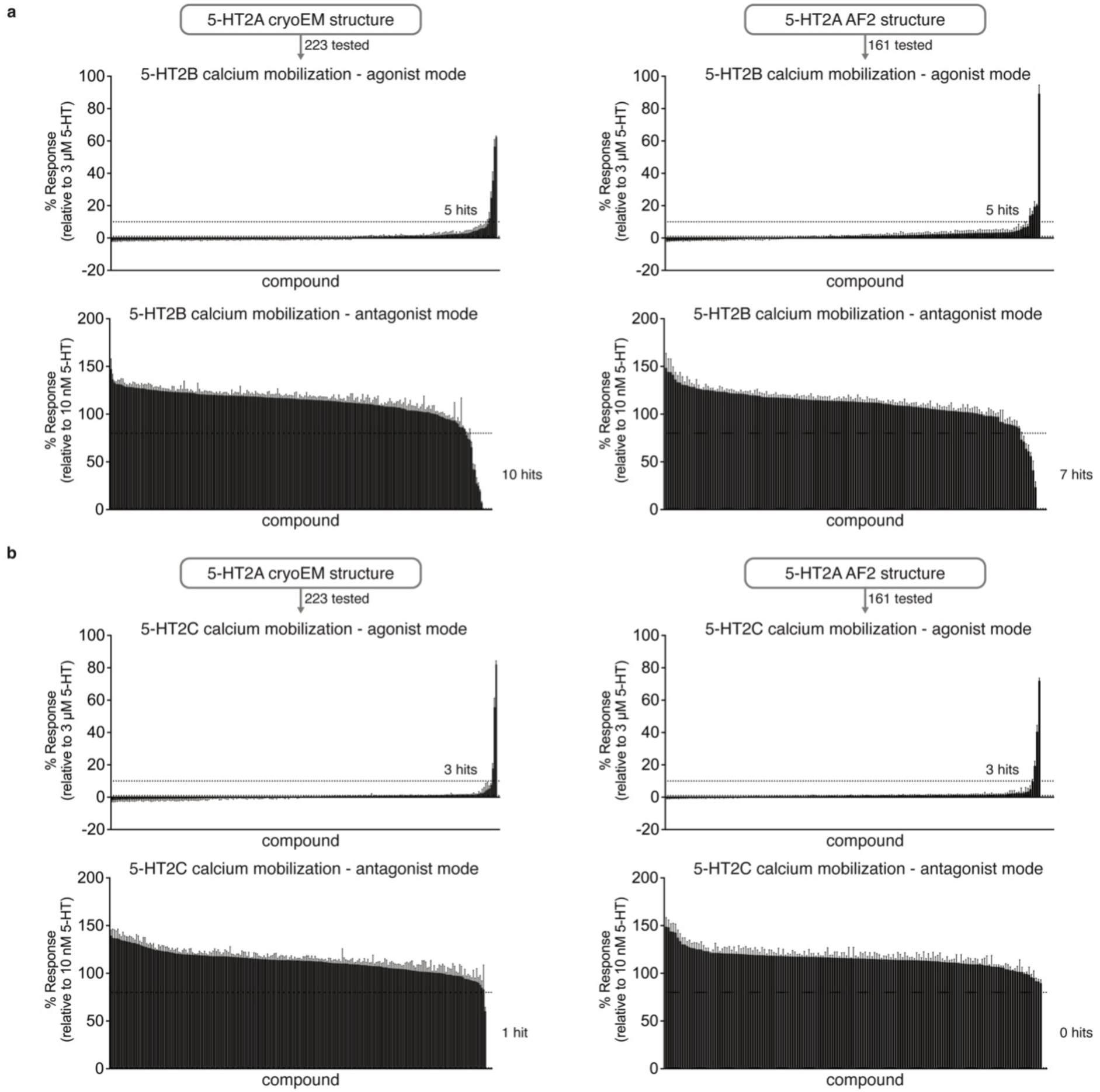
Functional screening of the prioritized library against the 5-HT2B and 5-HT2C receptors. Libraries from cryoEM (left) and AF2 (right) docking were screened at 3 μM in a calcium mobilization functional screen against the 5-HT2B (panel **a**) and 5-HT2C (panel **b**) receptors in agonist mode (top) or antagonist mode (bottom). Dashed lines indicate 10% maximal 5-HT response or 20% maximal clozapine response for agonists and antagonists, respectively. Data represent the mean ± s.e.m. from 3-4 biological replicates.

**Extended Data Fig. 10.**
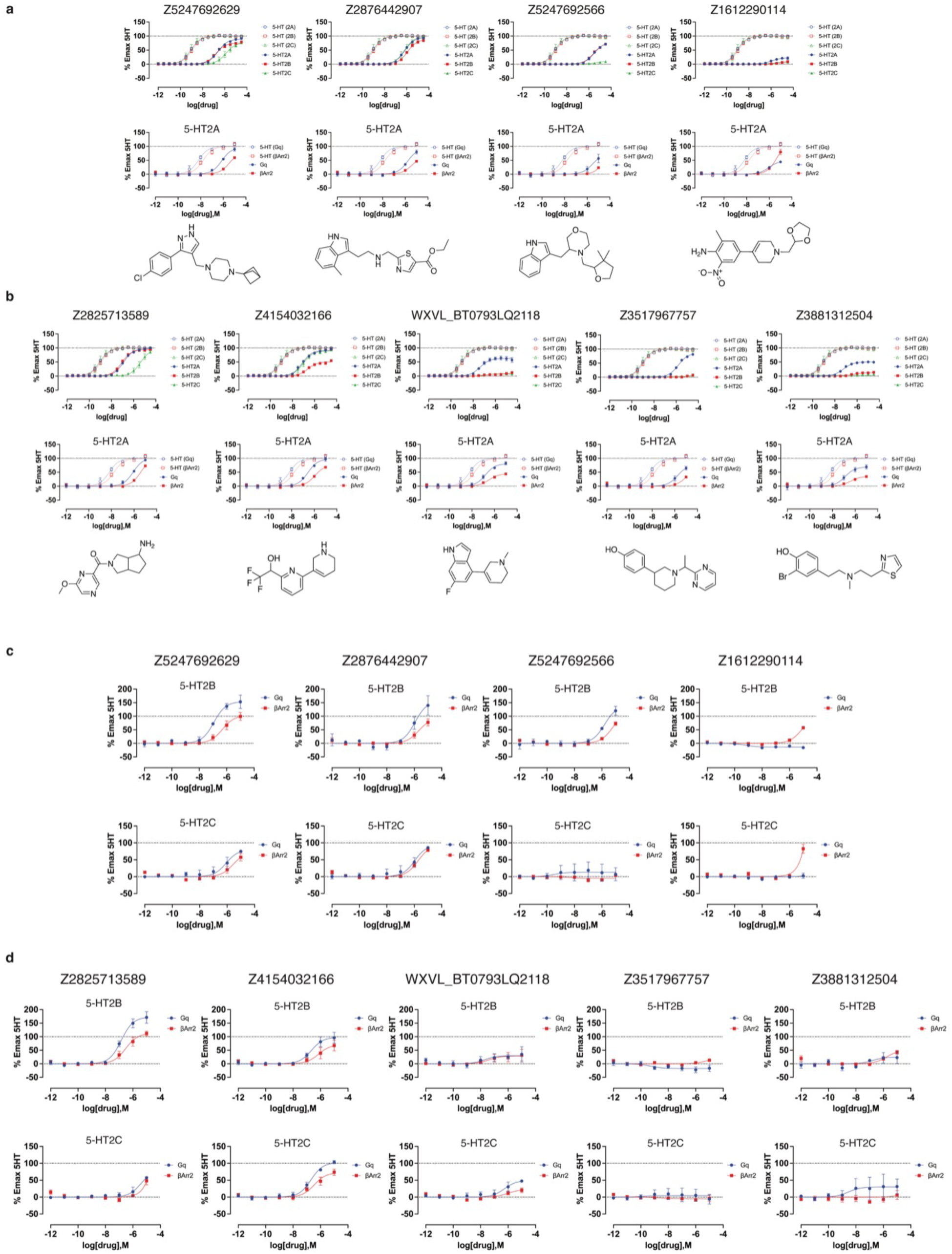
Signaling profiles of the top functional hits against the 5-HT2A, 5-HT2B, and 5-HT2C receptors. **a**. Functional assays for the top agonists from the cryoEM docking campaign relative to 5-HT (open symbols and dotted lines). Calcium mobilization (top) against 5-HT2A (blue), 5-HT2B (red), and 5-HT2C (green) receptors. BRET assays (bottom) measuring Gαq protein dissociation (blue) or β-arrestin2 recruitment (red) at the 5-HT2A receptor. **b**. Functional assays for the top agonists from the AF2 docking campaign relative to 5-HT (open symbols and dotted lines). Calcium mobilization (top) against 5-HT2A (blue), 5-HT2B (red), and 5-HT2C (green) receptors. BRET assays (bottom) measuring Gαq protein dissociation (blue) or β-arrestin2 recruitment (red) at the 5-HT2A receptor. The chemical structure of each compound is displayed below its respective dose-response curve. **c**. BRET assays measuring Gαq protein dissociation (blue) or β-arrestin2 recruitment (red) for the top agonists from the cryoEM docking campaign for 5-HT2B (top) and 5-HT2C (bottom) receptors. **d**. BRET assays measuring Gαq protein dissociation (blue) or β-arrestin2 recruitment (red) for the top agonists from the AF2 docking campaign for 5-HT2B (top) and 5-HT2C (bottom) receptors. Data are normalized relative to 5-HT and presented as mean ± s.e.m. of three biological replicates.

**Extended Data Fig. 11.**
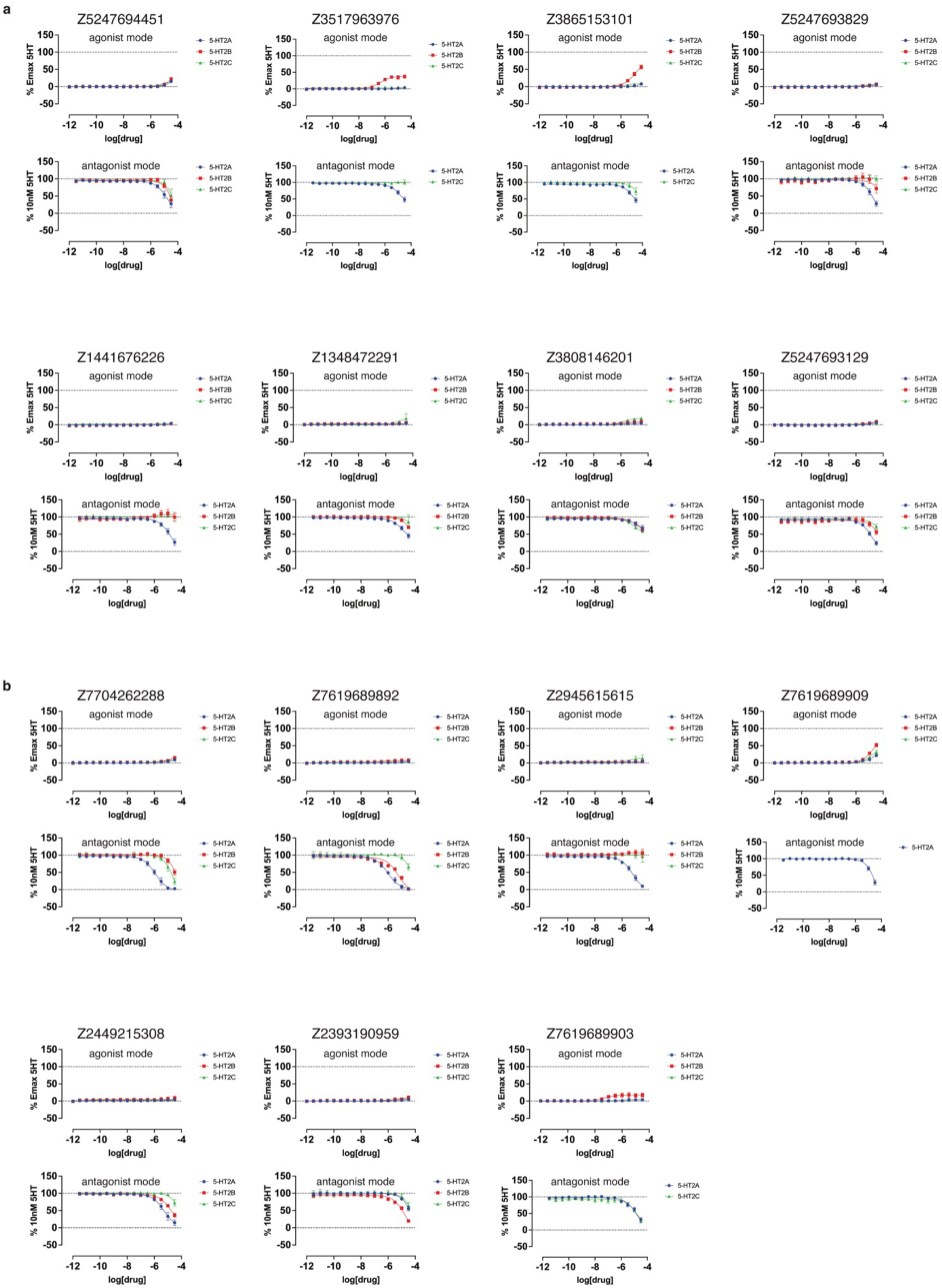
Functional dose-response curves measuring calcium mobilization of compounds that displace ≥ 90% [^3^H]-LSD at the 5-HT2A receptor. These compounds were tested at the 5-HT2A (blue), 5-HT2B (red), and 5-HT2C (green) receptors. **a.** Calcium mobilization assays performed in agonist mode (top) and antagonist mode (bottom) for compounds from the cryoEM docking set. **b.** Calcium mobilization assays performed in agonist mode (top) and antagonist mode (bottom) for compounds from the AF2 docking set. Data represent mean ± s.e.m. from three biological replicates.

**Extended Data Fig. 12.**
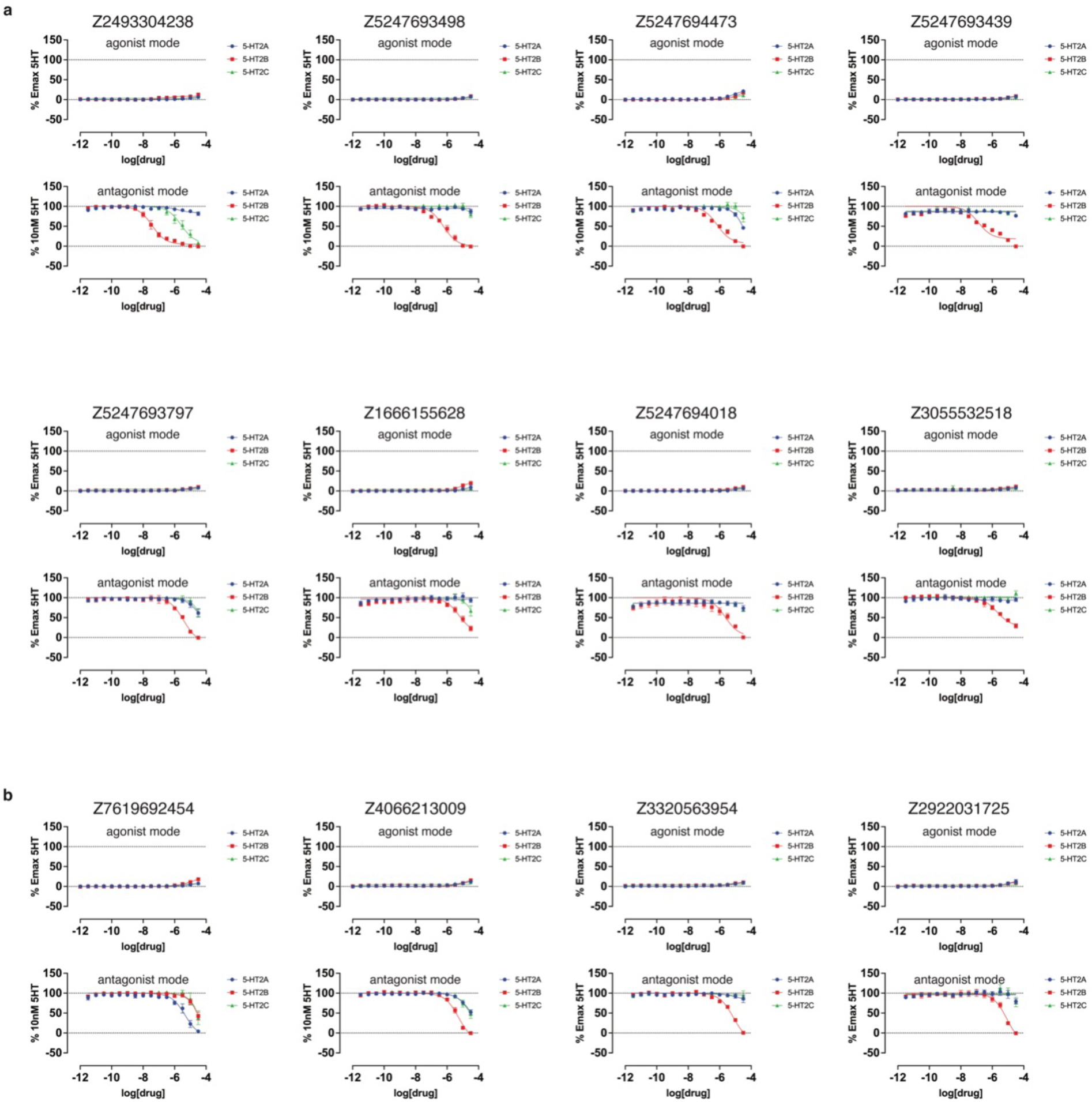
Functional dose-response curves measuring calcium mobilization of compounds exhibiting antagonist activity across any of the three 5-HT2-type receptors. These compounds were tested at the 5-HT2A (blue), 5-HT2B (red), and 5-HT2C (green) receptors. **a.** Calcium mobilization assays performed in agonist mode (top) and antagonist mode (bottom) for compounds from the cryoEM docking set. **b.** Calcium mobilization assays performed in agonist mode (top) and antagonist mode (bottom) for compounds from the AF2 docking set. Data represent mean ± s.e.m. from three biological replicates.

**Extended Data Fig. 13.**
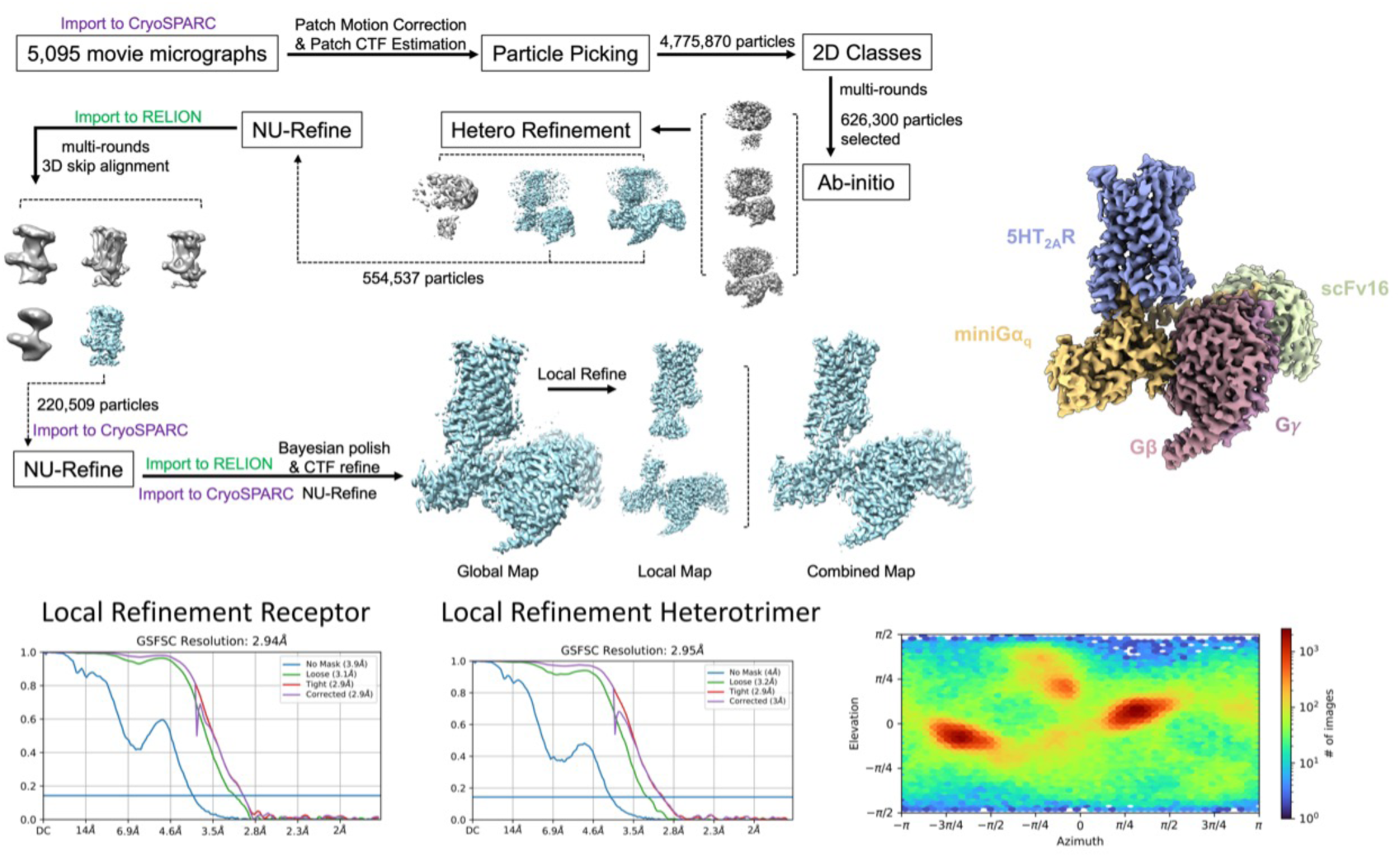
CryoEM Processing for Z7757. Comprehensive processing flow for the Z7757 structure. After 2D-classification and several rounds of 3D classification in cryoSPARC, the particles were transferred to for no-alignment 3D-classification focused on the receptor. Once a good consensus set of particles was identified and a further NU-refinemnet carried out, Bayesian polishing was performed on the particle set. A focused refinement was done on the receptor and a consensus map generated using Chimera. FSC plots for the receptor and heterotrimer as well as an angular distribution plot are also shown.

**Extended Data Fig. 14.**
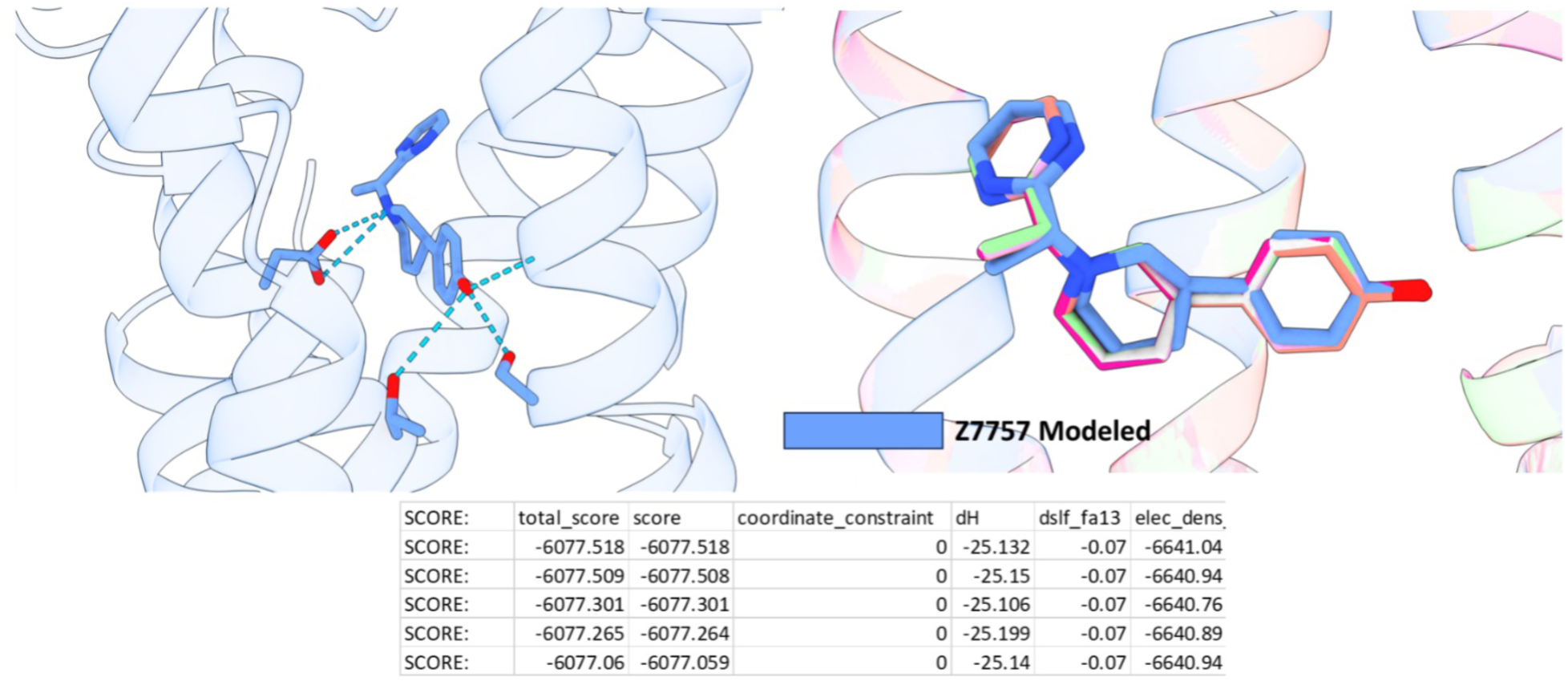
Emerald Docking recapitulates Z7757 binding pose. Shown in cornflower blue is the modeled binding pose of Z7757. The top 5 poses from Emerald, a docking algorithm that utilizes the cryoEM map as restraints, recapitulates the modeled binding pose. Below is a table of the top 5 scores output by Rosetta/Emerald.

**Extended Data Fig. 15.**
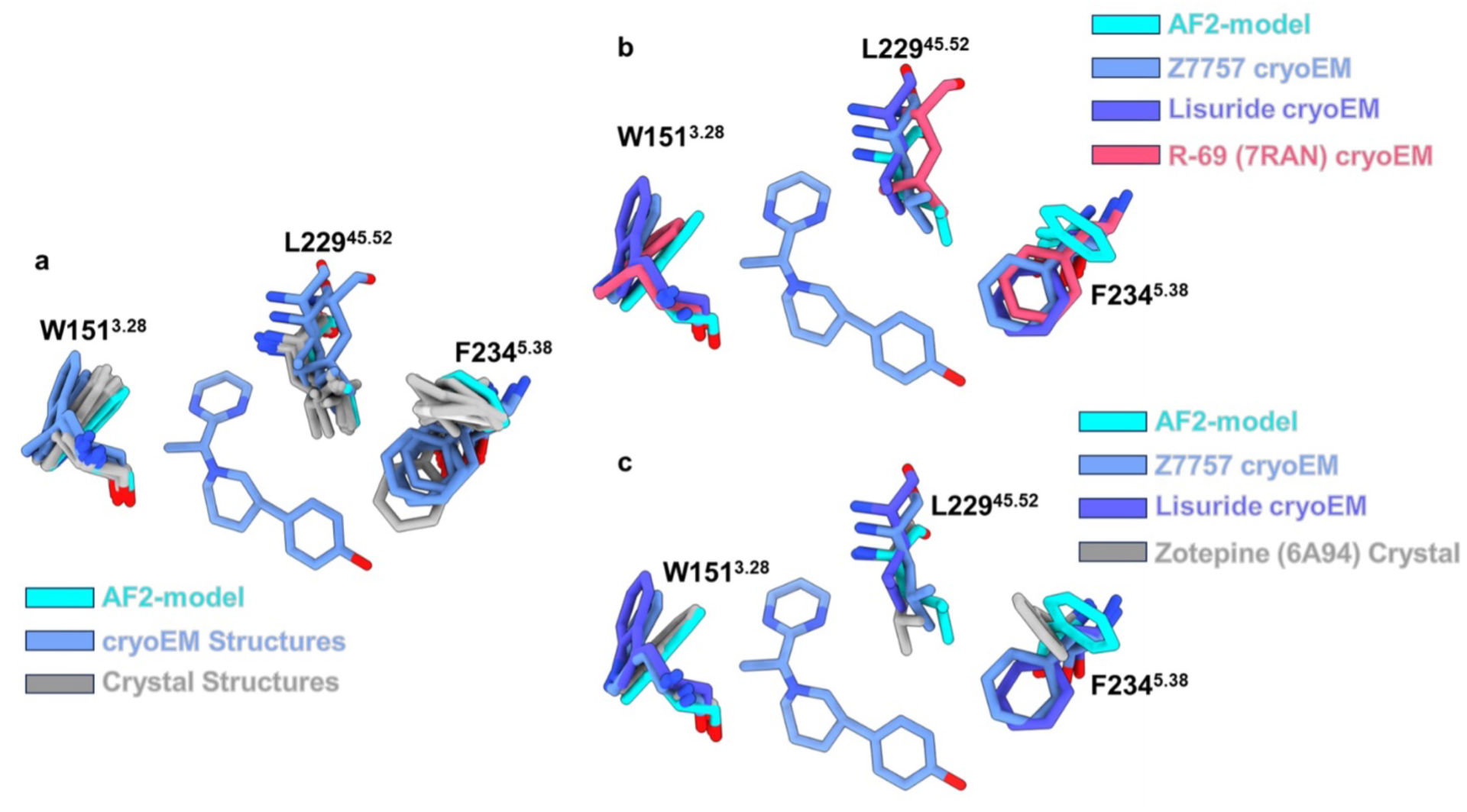
Alignment of all known 5-HT2A Structures. **(a)** All of the 5-HT2A structures were downloaded from the PDB and aligned via matchmaker in ChimeraX. Shown is the orthosteric pocket (and Z7757) with residues highlighted which showed the biggest differences between the two docking models. The structures are colored by method of determination AF2-model in cyan, cryoEM structures in cornflower blue, and crystal structures in grey. **(b)** Highlighted structural differences to show that a cryoEM structure can adopt the closed off pose of W151^3.28^ exhibited by the AF2-model. Here the AF2-model is shown in cyan, Z7757 cryoEM structure is shown in cornflower blue, the Lisuride cryoEM structure is shown in dark purple, and the agonist R-69 (PDB: 7RAN) cryoEM structure is shown in salmon. **(c)** Showing that an antagonist crystal structure can also adopt the closed position of W151^3.28^ exhibited by the AF2-model. Here the AF2-model is shown in cyan, Z7757 cryoEM structure is shown in cornflower blue, the Lisuride cryoEM structure is shown in dark purple, and the antagonist Zotepine (PDB: 6A94) cryoEM structure is shown in grey.

**Table S1.** CryoEM Data collection, model refinement, and validation.

| Structure | 5-HT2A-miniG<sub>q</sub>-Z7757 | 5-HT2A-miniG<sub>q</sub>-Lisuride |
| --- | --- | --- |
| PDB ID | 8V6U | 8UWL |
| **Data collection** |  |  |
| Magnification | 57,050 | 57,050 |
| Voltage (kV) | 300 | 300 |
| Dose per frame (e<sup>-</sup>/Å<sup>2</sup>) | 1.00 | 1.36 |
| Electron exposure (e<sup>-</sup>/Å<sup>2</sup>) | 50 | 67.78 |
| Defocus Range (μm) | -0.5 to -2.5 | -0.8 to -1.8 |
| Pixel size (Å) | 0.8677 | 0.8521 |
| Processing software | cryoSPARC/Relion | cryoSPARC |
| Symmetry imposed | C1 | C1 |
| Number of Micrographs | 5,095 | 3,282 |
| Initial particle images (no.) | 4,775,870 | 2,416,824 |
| Final particle images (no.) | 220,509 | 133,216 |
| Map resolution (Å) # | Local (2.9) / Global (3.0) | Receptor (3.1) / G protein (2.8) |
| FSC threshold | 0.143 | 0.143 |
| **Refinement Statistics\*** |  |  |
| **Model composition** |  |  |
| Number of chains | 5 | 5 |
| Total number of atoms | 8211 | 8626 |
| Number of Ligands | 1 | 1 |
| Number of residues | 1090 | 1123 |
| **Model validation** |  |  |
| CC (mask) map vs. model (%) | 0.77 | 0.83 |
| R.m.s. deviations |  |  |
| Bond lengths (Å) | 0.0004 | 0.013 |
| Bond angles (°) | 0.638 | 1.690 |
| Ramachandran plot |  |  |
| Favored (%) | 96 | 97 |
| Outlier (%) | 0 | 0 |
| Rotamer outliers (%) | 6.3 | 1.43 |
| Clash score | 6.47 | 5.41 |
| Molprobity score | 1.73 | 1.58 |

#Reported by cryoSPARC and \*Phenix comprehensive cryo-EM validation

**Table S2.** Binding affinities for compounds that displace ≥ 90% [^3^H]-LSD at the 5-HT2A receptor.

| Compound | log(Ki) | Ki | Ki (nM) | Model |
| --- | --- | --- | --- | --- |
| Z7704262288 | -7.82 ± 0.073 | 1.52E-08 | 15 | AF2 |
| Z7619689892 | -7.78 ± 0.077 | 1.68E-08 | 17 | AF2 |
| Z2945615615 | -7.62 ± 0.104 | 2.40E-08 | 24 | AF2 |
| Z5247694451 | -7.15 ± 0.073 | 7.10E-08 | 71 | cryoEM |
| Z7619689909 | -7.14 ± 0.078 | 7.22E-08 | 72 | AF2 |
| Z3881312504 | -7.02 ± 0.062 | 9.59E-08 | 96 | AF2 |
| Z3517963976 | -6.95 ± 0.085 | 1.13E-07 | 113 | cryoEM |
| Z3865153101 | -6.94 ± 0.099 | 1.14E-07 | 114 | cryoEM |
| Z2449215308 | -6.79 ± 0.100 | 1.62E-07 | 162 | AF2 |
| Z5247693829 | -6.77 ± 0.078 | 1.71E-07 | 171 | cryoEM |
| Z1441676226 | -6.77 ± 0.087 | 1.71E-07 | 171 | cryoEM |
| WXVL_BT0793LQ2118 | -6.68 ± 0.087 | 2.07E-07 | 207 | AF2 |
| Z1348472291 | -6.65 ± 0.083 | 2.22E-07 | 222 | cryoEM |
| Z3808146201 | -6.51 ± 0.096 | 3.08E-07 | 308 | cryoEM |
| Z2393190959 | -6.51 ± 0.085 | 3.12E-07 | 312 | AF2 |
| Z5247693129 | -6.49 ± 0.094 | 3.26E-07 | 326 | cryoEM |
| Z7619689903 | -6.46 ± 0.086 | 3.44E-07 | 344 | AF2 |

**Table S3.** Fit parameters from calcium mobilization and BRET assays for the top agonists from both docking sets against the 5-HT2A (top), 5-HT2B (middle), and 5-HT2C (bottom) receptor.

| Compound | 5-HT2A |  |  |  |  |  | Apparent Bias Factor | Apparent Bias for | Model |
| --- | --- | --- | --- | --- | --- | --- | --- | --- | --- |
|  | Calcium mobilization |  | Gaq dissociation |  | β-arrestin2 recruitment |  |  |  |  |
|  | pEC50 | Emax | pEC50 | Emax | pEC50 | Emax |  |  |  |
| 5-HT (reference) | 9.04 ± .0022 | 99.96 ± 0.64 | 8.18 ± 0.07 | 100.31 ± 2.45 | 7.78 ± 0.063 | 99.80 ± 2.34 | 1.0 | - | - |
| Z5247692629 | 6.61 ± 0.030 | 89.52 ± 1.21 | 6.15 ± 0.11 | 95.63 ± 5.59 | 5.48 ± 0.14 | 79.32 ± 8.00 | 2.2 | Gq | cryoEM |
| Z2876442907 | 6.19 ± 0.028 | 95.75 ± 1.37 | 5.80 ± 0.11 | 92.56 ± 6.63 | 5.41 ± 0.19 | 64.78 ± 9.55 | 1.4 | Gq | cryoEM |
| Z5247692566 | 5.52 ± 0.036 | 78.45 ± 1.87 | 5.53 ± 0.31 | 73.4 ± 16.73 | N.D. | N.D. | N.D. | N.D. | cryoEM |
| Z1612290114 | 5.94 ± 0.109 | 22.99 ± 1.37 | 6.03 ± 0.12 | 48.12 ± 3.22 | 5.31 ± 0.15 | 118.83 ± 15.27 | 0.8 | βarr2 | cryoEM |
| Z2825713589 | 6.92 ± 0.022 | 97.99 ± 0.88 | 6.08 ± 0.11 | 101.16 ± 5.99 | 5.40 ± 0.09 | 102.39 ± 7.42 | 1.8 | Gq | AF2 |
| Z4154032166 | 7.10 ± 0.029 | 91.96 ± 1.05 | 6.56 ± 0.11 | 97.68 ± 4.81 | 5.94 ± 0.07 | 75.77 ± 3.31 | 2.1 | Gq | AF2 |
| WXVL_BT0793LQ2118 | 7.38 ± 0.079 | 62.92 ± 1.86 | 6.81 ± 0.12 | 80.64 ± 4.13 | 6.50 ± 0.10 | 44.60 ± 2.06 | 1.4 | Gq | AF2 |
| Z3517967757 | 5.81 ± 0.031 | 85.64 ± 1.50 | 5.75 ± 0.15 | 76.30 ± 7.29 | 4.96 ± 0.67 | 69.34 ± 57.84 | 2.6 | Gq | AF2 |
| Z3881312504 | 7.23 ± 0.047 | 50.33 ± 0.92 | 6.89 ± 0.16 | 68.84 ± 4.82 | 6.41 ± 0.20 | 35.20 ± 3.39 | 2.3 | Gq | AF2 |

| Compound | 5-HT2B |  |  |  |  |  | Apparent Bias Factor | Apparent Bias for | Model |
| --- | --- | --- | --- | --- | --- | --- | --- | --- | --- |
|  | Calcium mobilization |  | Gαq dissociation |  | β-arrestin2 recruitment |  |  |  |  |
|  | pEC50 | E<sub>max</sub> | pEC50 | E<sub>max</sub> | pEC50 | E<sub>max</sub> |  |  |  |
| 5-HT (reference) | 8.95 ± 0.021 | 99.95 ± 0.64 | 9.19 ± 0.12 | 99.64 ± 3.52 | 8.96 ± 0.11 | 99.37 ± 3.55 | 1.0 | - | - |
| Z5247692629 | 6.80 ± 0.028 | 75.74 ± 0.91 | 6.94 ± 0.13 | 153.94 ± 8.71 | 6.32 ± 0.17 | 102.79 ± 8.95 | 3.6 | Gq | cryoEM |
| Z2876442907 | 5.92 ± 0.037 | 88.63 ± 1.82 | 5.95 ± 0.25 | 156.65 ± 23.55 | 5.68 ± 0.21 | 93.74 ± 13.54 | 1.8 | Gq | cryoEM |
| Z5247692566 | 5.58 ± 0.025 | 76.72 ± 1.24 | 5.82 ± 0.18 | 138.61 ± 15.80 | 5.23 ± 0.24 | 116.14 ± 26.32 | 2.7 | Gq | cryoEM |
| Z1612290114 | 5.02 ± 0.219 | 12.10 ± 2.36 | N.D. | N.D. | 5.09 ± 0.21 | 104.93 ± 23.59 | N.D. | N.D. | cryoEM |
| Z2825713589 | 7.09 ± 0.021 | 94.14 ± 0.79 | 6.85 ± 0.12 | 172.53 ± 8.9 | 6.53 ± 0.15 | 112.35 ± 7.87 | 1.9 | Gq | AF2 |
| Z4154032166 | 6.60 ± 0.06 | 50.86 ± 1.37 | 6.67 ± 0.21 | 98.24 ± 9.36 | 6.15 ± 0.31 | 71.07 ± 12.21 | 2.7 | Gq | AF2 |
| WXVL_BT0793LQ2118 | N.D. | N.D. | 7.71 ± 0.77 | 29.82 ± 8.45 | 7.41 ± 0.65 | 28.97 ± 7.20 | 1.2 | Gq | AF2 |
| Z3517967757 | N.D. | N.D. | N.D. | N.D. | N.D. | N.D. | N.D. | N.D. | AF2 |
| Z3881312504 | N.D. | N.D. | 6.78 ± 0.94 | 24.54 ± 10.48 | 5.77 ± 0.30 | 50.33 ± 10.01 | 2.9 | Gq | AF2 |

| Compound | 5-HT2C |  |  |  |  |  | Apparent Bias Factor | Apparent Bias for | Model |
| --- | --- | --- | --- | --- | --- | --- | --- | --- | --- |
|  | Calcium mobilization |  | Gαq dissociation |  | β-arrestin2 recruitment |  |  |  |  |
|  | pEC50 | Emax | pEC50 | Emax | pEC50 | Emax |  |  |  |
| 5-HT (reference) | 9.10 ± 0.041 | 99.80 ± 1.21 | 8.30 ± 0.07 | 100.40 ± 2.23 | 8.37 ± 0.14 | 99.21 ± 4.48 | 1.0 | - | - |
| Z5247692629 | 5.90 ± 0.083 | 76.61 ± 3.57 | 6.08 ± 0.26 | 80.03 ± 11.94 | 5.55 ± 0.30 | 73.81 ± 15.67 | 4.2 | Gq | cryoEM |
| Z2876442907 | 6.20 ± 0.054 | 98.70 ± 2.66 | 5.98 ± 0.22 | 94.28 ± 12.48 | 5.74 ± 0.13 | 93.02 ± 8.14 | 2.0 | Gq | cryoEM |
| Z5247692566 | 5.36 ± 0.331 | 10.26 ± 2.44 | N.D. | N.D. | N.D. | N.D. | N.D. | N.D. | cryoEM |
| Z1612290114 | N.D. | N.D. | N.D. | N.D. | N.D. | N.D. | N.D. | N.D. | cryoEM |
| Z2825713589 | 5.39 ± 0.09 | 96.11 ± 5.66 | 5.43 ± 0.29 | 78.47 ± 17.50 | N.D. | N.D. | N.D. | N.D. | AF2 |
| Z4154032166 | 7.02 ± 0.07 | 89.53 ± 2.38 | 6.7 ± 0.11 | 102.53 ± 4.96 | 6.46 ± 0.19 | 72.61 ± 6.64 | 2.8 | Gq | AF2 |
| WXVL_BT0793LQ2118 | N.D. | N.D. | 6.27 ± 0.33 | 49.37 ± 8.54 | 6.01 ± 0.48 | 22.49 ± 6.56 | 4.7 | Gq | AF2 |
| Z3517967757 | N.D. | N.D. | N.D. | N.D. | N.D. | N.D. | N.D. | N.D. | AF2 |
| Z3881312504 | N.D. | N.D. | 8.56 ± 1.19 | 29.95 ± 11.26 | N.D. | N.D. | N.D. | N.D. | AF2 |

**Table S4.** Fit parameters for compounds with antagonist activity at any of the 5-HT2-type receptors.

| Compound | 5-HT<sub>2A</sub> |  |  | 5-HT<sub>2B</sub> |  |  | 5-HT<sub>2C</sub> |  |  | Model |
| --- | --- | --- | --- | --- | --- | --- | --- | --- | --- | --- |
|  | pIC<sub>50</sub> | IC<sub>50</sub> | Potency (nM) | pIC<sub>50</sub> | IC<sub>50</sub> | Potency (nM) | pIC<sub>50</sub> | IC<sub>50</sub> | Potency (nM) |  |
| Z7619689892 | 6.04 ± 0.061 | 9.07E-07 | 907 | 5.26 ± 0.085 | 5.53E-06 | 5530 | N.D. | N.D. | N.D. | AF2 |
| Z7704262288 | 5.94 ± 0.065 | 1.16E-06 | 1157 | N.D. | N.D. | N.D. | 4.36 ± 0.261 | 4.34E-05 | 43421 | AF2 |
| Z7619692454 | 5.33 ± 0.103 | 4.63E-06 | 4635 | 3.95 ± 0.495 | 1.12E-04 | 111789 | N.D. | N.D. | N.D. | AF2 |
| Z2449215308 | 5.28 ± 0.091 | 5.20E-06 | 5198 | 4.81 ± 0.085 | 1.56E-05 | 15604 | N.D. | N.D. | N.D. | AF2 |
| Z2945615615 | 5.21 ± 0.072 | 6.23E-06 | 6234 | N.D. | N.D. | N.D. | N.D. | N.D. | N.D. | AF2 |
| Z1348472291 | 4.88 ± 0.116 | 1.31E-05 | 13106 | N.D. | N.D. | N.D. | N.D. | N.D. | N.D. | cryoEM |
| Z7619689903 | 4.88 ± 0.118 | 1.32E-05 | 13193 | N.D. | N.D. | N.D. | 4.64 ± 0.231 | 2.30E-05 | 23042 | AF2 |
| Z5247694451 | 4.81 ± 0.144 | 1.57E-05 | 15658 | N.D. | N.D. | N.D. | N.D. | N.D. | N.D. | cryoEM |
| Z3517963976 | 4.80 ± 0.133 | 1.58E-05 | 15773 | N.D. | N.D. | N.D. | N.D. | N.D. | N.D. | cryoEM |
| Z5247693129 | 4.80 ± 0.106 | 1.59E-05 | 15896 | N.D. | N.D. | N.D. | N.D. | N.D. | N.D. | cryoEM |
| Z1441676226 | 4.77 ± 0.142 | 1.69E-05 | 16924 | N.D. | N.D. | N.D. | N.D. | N.D. | N.D. | cryoEM |
| Z3808146201 | 4.72 ± 0.266 | 1.93E-05 | 19276 | 4.78 ± 0.178 | 1.65E-05 | 16453 | 5.07 ± .120 | 8.50E-06 | 8500 | cryoEM |
| Z5247693829 | 4.57 ± 0.168 | 2.66E-05 | 26645 | N.D. | N.D. | N.D. | N.D. | N.D. | N.D. | cryoEM |
| Z3865153101 | 4.57 ± 0.182 | 2.70E-05 | 26961 | N.D. | N.D. | N.D. | N.D. | N.D. | N.D. | cryoEM |
| Z4066213009 | 4.5 ± 0.157 | 3.17E-05 | 31687 | 5.27 ± 0.053 | 5.36E-06 | 5362 | 4.42 ± 0.263 | 3.82E-05 | 38237 | AF2 |
| Z5247694473 | 4.11 ± 0.519 | 7.82E-05 | 78210 | 6.16 ± 0.073 | 6.91E-07 | 691 | N.D. | N.D. | N.D. | cryoEM |
| Z7619689909 | 3.94 ± 0.384 | 1.14E-04 | 113676 | N.D. | N.D. | N.D. | N.D. | N.D. | N.D. | AF2 |
| Z2493304238 | N.D. | N.D. | N.D. | 7.43 ± 0.051 | 3.69E-08 | 37 | 5.58 ± 0.094 | 2.61E-06 | 2610 | cryoEM |
| Z5247693439 | N.D. | N.D. | N.D. | 6.87 ± 0.136 | 1.34E-07 | 134 | N.D. | N.D. | N.D. | cryoEM |
| Z5247693498 | N.D. | N.D. | N.D. | 6.23 ± 0.064 | 5.86E-07 | 586 | N.D. | N.D. | N.D. | cryoEM |
| Z3055532518 | N.D. | N.D. | N.D. | 5.69 ± 0.092 | 2.03E-06 | 2028 | N.D. | N.D. | N.D. | cryoEM |
| Z5247694018 | N.D. | N.D. | N.D. | 5.65 ± 0.181 | 2.25E-06 | 2248 | N.D. | N.D. | N.D. | cryoEM |
| Z5247693797 | N.D. | N.D. | N.D. | 5.43 ± 0.082 | 3.68E-06 | 3684 | N.D. | N.D. | N.D. | cryoEM |
| Z1666155628 | N.D. | N.D. | N.D. | 5.25 ± 0.168 | 5.68E-06 | 5684 | N.D. | N.D. | N.D. | cryoEM |
| Z3320563954 | N.D. | N.D. | N.D. | 5.14 ± 0.054 | 7.25E-06 | 7247 | N.D. | N.D. | N.D. | AF2 |
| Z2922031725 | N.D. | N.D. | N.D. | 5.06 ± 0.113 | 8.62E-06 | 8623 | N.D. | N.D. | N.D. | AF2 |
| Z2393190959 | N.D. | N.D. | N.D. | 4.77 ± 0.131 | 1.69E-05 | 16875 | N.D. | N.D. | N.D. | AF2 |

## Methods

### Molecular docking against the experimental structure of the σ_2_ and 5-HT2A receptors

The docking setups of the σ_2_ and 5-HT2A campaigns was reported previously^7,10^. The compound selection of the 5-HT2A campaign is described in the section below.

### Molecular docking against the AF2 structure of the σ_2_ and 5-HT2A receptors

The molecular docking procedures were conducted using the AF2 models of the σ_2_ and 5-HT2A receptors, specifically the AF-Q5BJF2-F1-model_v1 for the σ_2_ receptor and the AF-P28223-F1-model_v1 for the 5-HT2A receptor, both of which were downloaded from the AlphaFold Protein Structure Database (https://alphafold.ebi.ac.uk). The structures of both receptors were protonated at pH 7.0 using Epik and PROPKA in 2021 released Maestro. AMBER united atom charges were assigned to both structures. For the σ_2_ receptor, E73 was modeled as a neutral residue. The receptor was embedded in a lipid-bilayer to mimic its native environment in the endoplasmic reticulum (ER) membrane, followed by a 50ns coarse-grained molecular dynamic (MD) simulation with a restricted receptor conformation. The low dielectric and desolvation volumes were extended out 2.4 Å and 0.6 Å, respectively. For the 5-HT2A receptor, the low dielectric and desolvation volumes were extended out from the receptor surface by 2.0 Å and 0.6 Å, respectively. Energy grids for both receptors were pre-generated using CHEMGRID for AMBER van der Waals potential, QNIFFT for Poisson–Boltzmann-based electrostatic potentials, and SOLVMAP for ligand desolvation. The spheres generated by SPHGEN in the ligand binding site were used as matching spheres for docking known binders for each receptor against each apo AF2 model. Based on the predicted interactions from docking against apo AF2 models, the best docked pose of known binders for each receptor was then selected as matching spheres for the next steps. Twenty-seven spheres from the docked pose of PB28 were used for the σ_2_ receptor. Eighteen spheres of Lisuride were used for the 5-HT2A receptor. In order to speed up screening 1.6 billion molecules for the 5-HT2A receptor, the 5-HT2A matching spheres were labeled according to charge-charge interaction and hydrogen-bond patterns and grouped into four clusters via k-means clustering for efficient searching in docking calculations.

The docking setups for both the σ_2_ and 5-HT2A receptors were evaluated for their ability to enrich known ligands over property-matched decoys. The docking performance was evaluated using log-adjusted AUC values (logAUC). For the σ_2_ receptor, the enrichment achieved a logAUC of 16 on the same ligand-decoy set used to evaluate the docking setup built from the experimental structure. We used the same ‘extrema’ set to ensure that molecules with extreme physical properties were not enriched. The docking setup enriched close to 80% mono-cations among the top 1000 ranking molecules. The logAUC was 36 using ten σ_2_ ligands against the ‘extrema’ set. For the 5-HT2A receptor, this setup achieved a logAUC of 1.5 using the same ligand-decoy set as preparing the docking setup built from the cryoEM structure. On the extrema test, the docking setup enriched over 90% mono-cations among the top 1000 molecules, with a logAUC of 19. Additionally, a ‘goldilocks’ set from the DUDE-Z web server was used to further test the setup, resulting in a logAUC of 10. The evaluation above confirmed the favorable docking parameters for both receptors for launching ultra-large-scale docking campaigns.

In the large-scale docking campaigns, 490 million cations from ZINC15 (http://zinc20.docking.org) were docked against the σ_2_ receptor, and over 1.6 billion library molecules from ZINC20/ZINC22 (http://zinc20.docking.org and https://cartblanche22.docking.org) were docked against the 5-HT2A receptor using DOCK3.8. On average, 2,435 and 219 orientations were explored for the σ_2_ and 5-HT2A receptors, respectively. 180 and 340 conformations were averaged sampled for the σ_2_ and 5-HT2A receptors, respectively. The total calculation times were 115,653 hours for the σ_2_ receptor and 138,836 hours for the 5-HT2A receptors, respectively.

The σ_2_ receptor AF2 campaign was processed by the same protocol used to process the docking campaign against the σ_2_ receptor x-ray structure. The top-ranking 300,000 molecules were filtered for novelty using the ECFP4-based Tc against 2,232 σ1/_2_ ligands in ChEMBL (https://www.ebi.ac.uk/chembl/) and 574 σ_2_ ligands from S2RSLDB (http://www.researchdsf.unict.it/S2RSLDB). Molecules with Tc ≥ 0.35 were eliminated. The remaining 213,805 molecules were filtered by three criteria: (1) total torsion strain energy of less than 8 units; (2) maximum strain energy per torsion angle of less than 3 units; and (3) forms a salt bridge with D29. The last filter was implemented based on LUNA (https://github.com/keiserlab/LUNA). The remaining 57,662 molecules were clustered by the LUNA 1,024-length binary fingerprint of a Tc = 0.32, resulting in 12,095 clusters. Ultimately, 81 compounds were chosen by human inspection and 24 consecutive ranked compounds were selected from three rankings: the 1^st^, 550^th^, and 3925^th^, the same machine picking ranks as the docking campaign against the x-ray structure. In total, 153 compounds were selected by these two ways described above, 119 of which were successfully synthesized.

The 5-HT2A receptor campaigns against both experimental and AF2 structures were processed by the same protocol. The top-ranking 3.2 million molecules from both campaigns were filtered for novelty using the ECFP4-based Tc against 8,601 5-HT2A/2B/2C ligands in ChEMBL (https://www.ebi.ac.uk/chembl/). Molecules with Tc ≥ 0.35 were eliminated. The remaining top 2.67 and 2.64 million molecules were filtered by interactions with D155, S242, S239, T160, S159, and F340 for experimental and AF2 campaigns, respectively. The interaction filters were implemented based on LUNA (https://github.com/keiserlab/LUNA). The remaining 4,480 and 1,452 molecules were clustered by the LUNA 1,024-length binary fingerprint of a Tc = 0.35, resulting in 2,298 and 828 clusters. Ultimately, 259 and 218 compounds were chosen by human inspection. 223 and 161 compounds were successfully synthesized **(Supplementary Information Data 1)**.

### Competition binding in Expi293 membranes

Membranes were prepared as previously described^7^. Briefly, the human σ_2_ receptor was cloned into pcDNA3.1 (Invitrogen) mammalian expression vector with an amino-terminal protein C tag followed with a 3C protease cleavage site and transfected into Expi293 cells (Thermo Fisher Scientific) using FectoPRO (Polyplus-transfection) according to manufacturer instruction. Cells were harvested by centrifugation and lysed by osmotic shock in a buffer containing 20 mM HEPES, pH 7.5, 2 mM MgCl2,1:100,000 (vol/vol) benzonase nuclease (Sigma Aldrich), and cOmplete Mini EDTA-free protease-inhibitor tablets (Sigma Aldrich). The lysates were homogenized with a glass Dounce tissue homogenizer and then centrifuged at 20,000 x g for 20 min. Pellet was resuspended in 50 mM Tris, pH 8.0, adjusted to 10 mg/ml, flash frozen in liquid nitrogen in, and stored at 80 °C until use.

Competition binding reactions were done in 100 μL, with 50 mM Tris pH 8.0, 10 nM [^3^H]-DTG (PerkinElmer), indicated concentration of the competing ligand, and supplemented with 0.1% bovine serum albumin to minimize non-specific binding. Samples were shaken at 37 °C for 90 min. Afterward, the reaction was terminated by massive dilution and filtration over a glass microfiber filter with a Brandel harvester. Filters were soaked with 0.3% polyethyleneimine for at least 30 min before use. Radioactivity was measured by liquid scintillation counting. Data analysis was done in GraphPad Prism 9.0, with K_i_ values calculated by Cheng-Prusoff correction using the experimentally measured probe dissociation constant.

### Radioligand binding assays

Competitive radioligand binding assays were conducted in the NIMH PDSP using 5-HT2A membranes prepared from transiently transfected HEK293 cells in 96-well plates. Detailed assay protocols and conditions are also available from NIMH PDSP homepage (https://pdsp.unc.edu/pdspweb/?site=assays)

Compounds (10 mM DMSO stocks) were prepared in a standard binding buffer (50 mM Tris-HCl, pH 7.4, containing 10 mM MgCl_2_ and 0.1 mM EDTA) supplemented with 1 mg/mL fatty-acid free bovine serum albumin and 0.1 mg/mL ascorbic acid. Samples were first screened at a single concentration of 10 μM with in-plate quadruplicate set (primary binding experiment). Those with a minimum of 90% inhibition at 10 μM were subjected to concentration response assays (12 points starting from 100 μM) to determine binding affinity (Ki) with in-plate duplicate set for 3 independent assays (secondary binding experiment). Radioligand [3H]-LSD were used at the final concentration of 0.5 nM. To each well of a 96-well plate, containing 25 μL of the test compounds at 50 μM, 25 μL of the radioligand was added, followed by the addition of 75 μL of crude membrane fractions containing the 5-HT2A receptor in standard binding buffer. For primary binding experiments, nonspecific binding was determined in the presence of 10 μM Clozapine. The reaction mixture was incubated at room temperature for 2 hours. Receptor-bound radioactivity was harvested by rapid filtration onto UniFilter-96 GF/C Microplates (PerkinElmer) using a Filtermate harvester (PerkinElmer). The dried plates were treated with MicroScint-O liquid scintillation cocktails (PerkinElmer), and the captured radioactivity was measured using a MicroBeta scintillation counter. Concentration response inhibition results were analyzed utilizing the “Binding Competitive – One site Fit Ki” model in GraphPad Prism version 10.1.0 (GraphPad Prism).

### Calcium Mobilization Assay

Calcium mobilization was measured using Fluo-4 Direct calcium dye (Invitrogen). HEK293 cell lines stably transfected with 5-HT2A, 5-HT2B, or 5-HT2C were maintained in Dulbecco’s Modified Eagle Media (DMEM) supplemented with 10% fetal bovine serum (FBS), 100 I.U./mL penicillin, 100 mg/mL streptomycin, 100 μg/mL hygromycin B, and 15 μg/mL blasticidin in a humidified incubator at 37°C and 5% CO_2_. To induce receptor expression, cells were treated with 1.5 μg/mL tetracycline and seeded into 384-well black plates in DMEM supplemented with 1% dialyzed FBS (dFBS), 100 I.U./mL penicillin, and 100 mg/mL streptomycin at a cell density of ∼15,000 cells/well. 24 hours later, cells were incubated with 20 μL Fluo-4 Direct calcium dye (Invitrogen) supplemented with 2 mM probenecid (Thermofisher) in drug buffer (1X HBSS, 20 mM HEPES, 0.1% (w/v) BSA, 0.01% (w/v) ascorbic acid, pH = 7.4) for 1 hour at 37°C and 5% CO_2_, then 20 minutes at room temperature in the dark. All drug dilutions were prepared at a 3X concentration in drug buffer for screening (3 μM final concentration) or dose-response profiling (16-point titration) and transferred into a 384-well drug plate. Calcium flux was quantified using a FLIPR^PENTA^ fluorescence imaging plate reader (Molecular Dynamics) by first measuring baseline for 10 seconds (1 read per second), then following addition of 10 μL 3X drug for 120 seconds (1 read per second). For antagonist data, 10 μL 5-HT (10 nM final concentration) was added 15 minutes after drug treatment to stimulate calcium mobilization. Maximal raw fluorescence values over the 120-second experiment were transformed to fold-change relative to baseline fluorescence (mean of the first 10 reads prior to drug addition) and plotted for each dose to generate endpoint screening data or dose-response curves. These data were normalized relative to serotonin (agonist control) or clozapine (antagonist control) and plotted as bar graphs (screening data) or fit using the “log(agonist) vs. response (three parameter)” or “log(inhibitor) vs. response (three parameter)” functions in GraphPad Prism version 10.1.0 (GraphPad Software). An activity threshold of 10% 5-HT response or 20% clozapine response was used to identify compounds that exhibit agonist or antagonist activity, respectively. Importantly, dose-response curves for compounds with agonist activity (≥ 10% maximal 5-HT response at 10 μΜ) at a given receptor were removed from antagonist mode plots.

### Bioluminescence resonance energy transfer assays (BRET)

Bioluminescence resonance energy transfer (BRET) assays were performed to measure Gα_q_ protein dissociation (BRET2) and β-arr2 recruitment (BRET1) as described previously (Olson 2020, Wang et al 2023). For Gα_q_ protein dissociation measurements, wildtype 5-HT2A, 5-HT2B and 5-HT2C were co-transfected with Gα_q_-RLuc8, G_β3_, and GFP2-G_γ9_ in a 1:1:1:1 ratio in HEK293 cells maintained in DMEM supplement with 10% FBS and 1% penicillin-streptomycin (pen-strep). After 12 h, cells were plated in 96-well microplates in DMEM supplemented with 1% dFBS and 1% pen-strep. After 12 h of plating the cells, media was vacuum aspirated and followed by addition of 60 μL of assay buffer (1x Hank’s Balanced Salt Solution in phosphate buffered saline, 20 mM HEPES, pH 7.4) into the wells and plates were incubated at 37 °C for 10 min. After that 30 μL of 3x drug diluted in drug buffer (assay buffer supplemented with 0.1% fatty acid-free bovine serum albumin and 0.01 % ascorbic acid) were added to the wells. Plates were incubated at 37 °C for 10 min. Plates were then removed and kept at RT for another 10 min followed by addition of 10 μL of coelenterazine 400a diluted in assay buffer at a concentration of 50 μM. Plates were incubated for another 10 min at RT and then read using a BMG Labtech PHERAstar FSX with BRET2 plus optic module. For BRET1, RLuc8 was cloned directly to the C-terminus of wildtype 5-HT2A, 5-HT2B and 5-HT2C and experiments were performed as described previously (wang et al 2023). HEK293T cells were co-transfected with receptor, GRK2 and mVenus-β-arr2 in a 1:1:5 ratio. BRET1 measurements were done identically to BRET2, though coelenterazine h was used instead of coelenterazine 400a. Plates were read using a BMG Labtech PHERAstar FSX with BRET1 plus optic module. All data were analyzed using GraphPad Prism 9.0.

### Protein Purification

The protein purification and complex formation was carried out as previously described^54,62,64^. After expression in *Spodoptera frugiperda* (Sf9) the 5HT2A receptor (with the respective ligand), mini-Gαq, and scFv16 were purified as previously described. In the case of the Lisuride complex, each of the component was mixed in a 1:1.2:1.5 molar equivalent ratio of receptor:mini-Gαq:scFv16 and allowed to incubate overnight at 4°C. The mixture was further purified by Superdex 200 10/300 column in 20 mM HEPES (pH 7.5), 100 mM NaCl, 0.001% (w/v) LMNG, 0.0001% (w/v) CHS, 0.00025% (w/v) GDN, 100 μM TCEP, 50 μM Lisuride. For the Z7757 structure, the 5HT2A receptor and mini-Gαq heterotrimer were co-expressed utilizing an MOI of 3:1.5, respectively. Isolation of the complex was carried out in the same way as the receptor purification, with the addition of 500 μg of scFv16 during membrane solubilization. The first round of size exclusion was carried out in 20 mM HEPES (pH 7.5), 100 mM NaCl, 0.001% (w/v) LMNG, 0.0001% (w/v) CHS, 0.00025% (w/v) GDN, 100 μM TCEP, 50 μM Z7757 on a Superose 6 column and peak fractions corresponding to the complex were collected. Each sample was concentrated to 3mg/mL for the Z7757 sample or 15 mg/mL for the Lisuride sample prior to preparing grids.

### cryoEM Data Collection and Processing

Grids were prepared by applying 3.5 μLs of the Z7757 complex or 2 μLs of the Lisuride complex to glow discharged UltrAuFoil holey gold grids (Quantifoil, Au300-R1.2/1.3). The girds containing Z7757 were then vitrified by plunge freezing into a 60/40 mixure of liquid ethane/propane using a Vitrobot mark IV (FEI) set at 22°C and 100% humidity while the grids containing Lisuride were vitrified by plunge freezing into liquid ethane at 18°C and 100% humidity. Movies were then collected on a Titan Krios operated at 300 keV with a K3 Summit direct electron detector (Gatan) at a magnification of 57,050x. A total of 5,095 movies were collected with 50 frames per movie over a total dose of 50 electrons/Å2 and a defocus range 0.5μm to 2.5μm for Z7757. For the lisuride complex, a total of 3,282 movies were collected, dose-fractionated over 50 frames, and recorded for 0.05 sec/frame, resulting in a total dose of 67.78 electrons/Å² in super-resolution mode with a defocus range of 0.8-1.8 μm. The data processing tree can be found for the Z7757 dataset in **Extended Data Figure 13.** In short, motion correction and CTF estimation was done using cryoSPARC ^67^ and initial sets of particle picking, 2D classification, initial models, and 3D refinement. A set of approximately 554,537 particles were then carried over into Relion ^68^ for multiple rounds of no-alignment 3D classification focused on the TM domain. Subsequent rounds of NU-Refine (carried out in cryoSPARC) and Bayesian polishing (carried out in Relion) were then done. Once a good consensus map was achieved from NU-refine, subsequent local refinement with a soft mask was carried out in cryosparc and the global/local maps were combined. For the lisuride complex, the processing tree can be found in **Extended Data Figure 5.** In short, the data processing was done in cryosparc with a total of 2,426,824 particles extracted from the corrected micrographs. Following 2D and 3D classification, a subset of 133,216 particles underwent homogeneous refinement, followed by local refinements of the Lisuride-bound receptor and the heterotrimeric miniGq (bound to the active-state stabilizing single-chain variable fragment, scFv16) at resolutions of 3.1 Å and 2.8 Å, respectively. Maps resulting from local refinements were combined in Chimera to produce the final reconstruction.

### Model Building and Refinement

The initial model was fit using PDB 7RAN ^49^. All models were docked into the cryoEM density using either Chimera ^69^ or ChimeraX ^70^. Subsequent real-space refinement was carried out in phenix^71^ and iterative manual fitting and adjustments were carried out with COOT ^72^. The binding pose was further validated by using Emerald for Z7757 and the Gemspot pipeline for Lisuride ^63^ ^55^. Final model statistics were validated by Molprobity^73^ (**Extended Data Table 1**). All structural figures were visualized either in Chimera^69^ or ChimeraX^70^.

## Acknowledgements

Funding was provided by US NIH grants R35GM122481 (to BKS), by GM71896 (to JJI), by US NIH grants RO1MH112205, R37DA045657 and DARPA (to BLR) and R01GM119185, the Vallee Foundation, and the Sanofi iAwards program (to ACK). We thank OpenEye Software for the use of Omega and Schrodinger LLC for the use of prepwizard in Maestro.

## Author Contributions

J.L., B.K.S., A.C.K. and B.L.R. conceived the study. J.L. conducted the docking, chemoinformatics analyses and ligand picking, assisted in the latter by T.A.T., S,H., and B.K.S.. N.K., M.K.J., K.S., Y.K., J.B. and B.L.R. performed the 5-HT2A/2B/2C receptor assays and analysis. R.G., L.W., X.B.A., K.K. G.K. and B.L.R. determined cryoEM 5-HT2A structures. A.A. and A.C.K. performed the σ_2_ receptor assays and analysis. O.O.T. and Y.S.M. supervised the synthesis of molecules from the virtual library. J.J.I. was responsible for the building of the version of the ZINC library that was docked. A.C.K., B.K.S. and B.L.R. supervised the project. The manuscript was written by J.L., B.K.S. N.K., R.G., and B.L.R. with input from other authors.

## Data availability

The compounds docked in this study are freely available from our ZINC20/22 database, http://zinc20.docking.org and https://cartblanche22.docking.org. The cryo-EM density map and corresponding coordinates for 5-HT2AR/miniGq/Lisuride and 5-HT2AR/miniGq/Z77575 have been deposited in the Electron Microscopy Data Bank (EMDB) and the Protein Data Bank (PDB). The 5-HT2AR/miniGq/Lisuride structure has been deposited under accession codes EMD-42676 and PDB ID: 8UWL, while the 5-HT2AR/miniGq/Z77575 structure EMD-42999 and PDB ID: 8V6U.

## Code availability

DOCK3.8 is freely available for non-commercial research http://dock.compbio.ucsf.edu/DOCK3.8/. A web-based version available to all is available at http://blaster.docking.org/.

## References

1 Sadybekov, A. V. & Katritch, V. Computational approaches streamlining drug discovery. Nature 616, 673–685 (2023).

2 Wang, Y. et al. Structures of the entire human opioid receptor family. Cell 186, 413–427. e417 (2023).

3 Xu, P. et al. Structural insights into the lipid and ligand regulation of serotonin receptors. Nature 592, 469–473 (2021).

4 Lyu, J. et al. Ultra-large library docking for discovering new chemotypes. Nature 566, 224–229 (2019).

5 Stein, R. M. et al. Virtual discovery of melatonin receptor ligands to modulate circadian rhythms. Nature 579, 609–614 (2020).

6 Gorgulla, C. et al. An open-source drug discovery platform enables ultra-large virtual screens. Nature 580, 663–668 (2020).

7 Alon, A. et al. Structures of the σ2 receptor enable docking for bioactive ligand discovery. Nature 600, 759–764 (2021).

8 Fink, E. A. et al. Structure-based discovery of nonopioid analgesics acting through the α2A-adrenergic receptor. Science 377, eabn7065 (2022).

9 Sadybekov, A. A. et al. Synthon-based ligand discovery in virtual libraries of over 11 billion compounds. Nature 601, 452–459 (2022).

10 Lyu, J., Irwin, J. J. & Shoichet, B. K. Modeling the expansion of virtual screening libraries. Nature Chemical Biology, 1–7 (2023).

11 Manglik, A. et al. Structure-based discovery of opioid analgesics with reduced side effects. Nature 537, 185–190 (2016).

12 Reis, J. et al. Targeting ROS production through inhibition of NADPH oxidases. Nature Chemical Biology, 1–11 (2023).

13 Gorgulla, C. et al. VirtualFlow 2.0-The Next Generation Drug Discovery Platform Enabling Adaptive Screens of 69 Billion Molecules. bioRxiv, 2023.2004. 2025.537981 (2023).

14 Hauser, A. S. et al. Pharmacogenomics of GPCR drug targets. Cell 172, 41–54. e19 (2018).

15 Carlsson, J. et al. Ligand discovery from a dopamine D3 receptor homology model and crystal structure. Nature chemical biology 7, 769–778 (2011).

16 Beuming, T. & Sherman, W. Current Assessment of Docking Into GPCR Crystal Structures and Homology Models: Successes, Challenges, and Guidelines. Journal of Chemical Information and Modeling, doi:10.1021/ci300411b (2012).

17 Hillisch, A., Pineda, L. A. & Hilgenfeld, R. Utility of Homology Models in the Drug Discovery Process. Drug Discovery Today, doi:10.1016/s1359-6446(04)03196-4 (2004).

18 Bordogna, A., Pandini, A. & Bonati, L. Predicting the accuracy of protein–ligand docking on homology models. Journal of computational chemistry 32, 81–98 (2011).

19 Jumper, J. et al. Highly accurate protein structure prediction with AlphaFold. Nature 596, 583–589 (2021).

20 Baek, M. et al. Accurate Prediction of Protein Structures and Interactions Using a Three-Track Neural Network. Science, doi:10.1126/science.abj8754 (2021).

21 Varadi, M. et al. AlphaFold Protein Structure Database: massively expanding the structural coverage of protein-sequence space with high-accuracy models. Nucleic acids research 50, D439–D444 (2022).

22 Tunyasuvunakool, K. et al. Highly Accurate Protein Structure Prediction for the Human Proteome. Nature, doi:10.1038/s41586-021-03828-1 (2021).

23 Hutin, S. et al. The Vaccinia virus DNA helicase structure from combined single-particle cryo-electron microscopy and AlphaFold2 prediction. Viruses 14, 2206 (2022).

24 Mosalaganti, S. et al. AI-based structure prediction empowers integrative structural analysis of human nuclear pores. Science 376, eabm9506 (2022).

25 Akdel, M. et al. A structural biology community assessment of AlphaFold2 applications. Nature Structural & Molecular Biology 29, 1056–1067 (2022).

26 Jendrusch, M., Korbel, J. O. & Sadiq, S. K. AlphaDesign: A de novo protein design framework based on AlphaFold. Biorxiv, 2021.2010. 2011.463937 (2021).

27 Goverde, C. A., Wolf, B., Khakzad, H., Rosset, S. & Correia, B. E. De novo protein design by inversion of the AlphaFold structure prediction network. Protein Science 32, e4653 (2023).

28 Bryant, P., Pozzati, G. & Elofsson, A. Improved prediction of protein-protein interactions using AlphaFold2. Nature communications 13, 1265 (2022).

29 Yin, R., Feng, B. Y., Varshney, A. & Pierce, B. G. Benchmarking AlphaFold for protein complex modeling reveals accuracy determinants. Protein Science 31, e4379 (2022).

30 Evans, R. et al. Protein complex prediction with AlphaFold-Multimer. biorxiv, 2021.2010. 2004.463034 (2021).

31 Wang, S. et al. CavitySpace: a database of potential ligand binding sites in the human proteome. Biomolecules 12, 967 (2022).

32 van Kempen, M. et al. Fast and accurate protein structure search with Foldseek. Nature Biotechnology, 1–4 (2023).

33 Ma, W. et al. Enhancing protein function prediction performance by utilizing AlphaFold-predicted protein structures. Journal of Chemical Information and Modeling 62, 4008–4017 (2022).

34 Hekkelman, M. L., de Vries, I., Joosten, R. P. & Perrakis, A. AlphaFill: enriching AlphaFold models with ligands and cofactors. Nature Methods 20, 205–213 (2023).

35 Weghoff, M. C., Bertsch, J. & Müller, V. A novel mode of lactate metabolism in strictly anaerobic bacteria. Environmental microbiology 17, 670–677 (2015).

36. Kimura, S., et al. Sequential action of a tRNA base editor in conversion of cytidine to pseudouridine. Nature Communications 13, 5994 (2022).

37 Tao, H. et al. Discovery of non-squalene triterpenes. Nature 606, 414–419 (2022).

38 Thornton, J. M., Laskowski, R. A. & Borkakoti, N. AlphaFold Heralds a Data-Driven Revolution in Biology and Medicine. Nature Medicine, doi:10.1038/s41591-021-01533-0 (2021).

39 Holcomb, M., Chang, Y.-T., Goodsell, D. S. & Forli, S. Evaluation Of *AlphaFold2* Structures as Docking Targets. Protein Science, doi:10.1002/pro.4530 (2022).

40 Karelina, M., Noh, J. J. & Dror, R. O. How accurately can one predict drug binding modes using AlphaFold models? bioRxiv, 2023.2005. 2018.541346 (2023).

41 He, X. et al. AlphaFold2 Versus Experimental Structures: Evaluation on G Protein-Coupled Receptors. Acta Pharmacologica Sinica, doi:10.1038/s41401-022-00938-y (2022).

42 Filippo, J. I. D. & Cavasotto, C. N. How Good Are AlphaFold Models for Docking-Based Virtual Screening?, doi:10.26434/chemrxiv-2022-sgj8c (2022).

43 Díaz-Rovira, A. M. & Guallar, V. Are Deep Learning Structural Models Sufficiently Accurate for Virtual Screening? Application of Docking Algorithms to AlphaFold2 Predicted Structures. Journal of Chemical Information and Modeling, doi:10.1021/acs.jcim.2c01270 (2023).

44. Wong, F. et al. Benchmarking *AlphaFold*-enabled Molecular Docking Predictions for Antibiotic Discovery. Molecular Systems Biology doi:10.15252/msb.202211081 (2022).

45 Zhang, Y. et al. Benchmarking Refined and Unrefined AlphaFold2 Structures for Hit Discovery. doi:10.26434/chemrxiv-2022-kcn0d-v2 (2022).

46 Díaz-Rovira, A. M. & Guallar, V. Are Deep Learning Structural Models Sufficiently Accurate for Free-Energy Calculations? Application of FEP+ to AlphaFold2-Predicted Structures. Journal of Chemical Information and Modeling, doi:10.1021/acs.jcim.2c00796 (2022).

47 Masha, K., Noh, J. & Dror, R. How accurately can one predict drug binding modes using AlphaFold models. Elife 12 (2023).

48. Lowe, D. in *IN THE PIPELINE* (2023).

49 Kaplan, A. L. et al. Bespoke library docking for 5-HT2A receptor agonists with antidepressant activity. Nature 610, 582–591 (2022).

50 Koehl, A. et al. Structure of the micro-opioid receptor-G(i) protein complex. Nature 558, 547–552, doi:10.1038/s41586-018-0219-7 (2018).

51 Xia, R. et al. Cryo-EM structure of the human histamine H(1) receptor/G(q) complex. Nat Commun 12, 2086, doi:10.1038/s41467-021-22427-2 (2021).

52 Mobbs, J. I. et al. Structures of the human cholecystokinin 1 (CCK1) receptor bound to Gs and Gq mimetic proteins provide insight into mechanisms of G protein selectivity. PLoS Biol 19, e3001295, doi:10.1371/journal.pbio.3001295 (2021).

53 Cao, C. et al. Signaling snapshots of a serotonin receptor activated by the prototypical psychedelic LSD. Neuron 110, 3154–3167 e3157, doi:10.1016/j.neuron.2022.08.006 (2022).

54 Gumpper, R. H., Fay, J. F. & Roth, B. L. Molecular insights into the regulation of constitutive activity by RNA editing of 5HT(2C) serotonin receptors. Cell Rep 40, 111211, doi:10.1016/j.celrep.2022.111211 (2022).

55 Robertson, M. J., van Zundert, G. C. P., Borrelli, K. & Skiniotis, G. GemSpot: A Pipeline for Robust Modeling of Ligands into Cryo-EM Maps. Structure 28, 707–716 e703, doi:10.1016/j.str.2020.04.018 (2020).

56 Cao, D. et al. Structure-based discovery of nonhallucinogenic psychedelic analogs. Science 375, 403–411 (2022).

57 Pándy-Szekeres, G. et al. GPCRdb in 2023: state-specific structure models using AlphaFold2 and new ligand resources. Nucleic Acids Research 51, D395–D402 (2023).

58 Isberg, V. et al. Generic GPCR residue numbers–aligning topology maps while minding the gaps. Trends in pharmacological sciences 36, 22–31 (2015).

59 Simon, I. A. et al. Ligand selectivity hotspots in serotonin GPCRs. Trends in Pharmacological Sciences.

60 Olsen, R. H. et al. TRUPATH, an open-source biosensor platform for interrogating the GPCR transducerome. Nature chemical biology 16, 841–849 (2020).

61 Liu, Y. et al. Ligand recognition and allosteric modulation of the human MRGPRX1 receptor. Nat Chem Biol 19, 416–422, doi:10.1038/s41589-022-01173-6 (2023).

62 Cao, C. et al. Structure, function and pharmacology of human itch GPCRs. Nature 600, 170–175, doi:10.1038/s41586-021-04126-6 (2021).

63 Muenks, A., Zepeda, S., Zhou, G., Veesler, D. & DiMaio, F. Automatic and accurate ligand structure determination guided by cryo-electron microscopy maps. Nature Communications 14, 1164 (2023).

64 Kim, K. et al. Structure of a hallucinogen-activated Gq-coupled 5-HT2A serotonin receptor. Cell 182, 1574–1588. e1519 (2020).

65 Krishna, R. et al. Generalized Biomolecular Modeling and Design with RoseTTAFold All-Atom. bioRxiv, 2023.2010. 2009.561603 (2023).

66. Google DeepMind AlphaFold Team, Isomorphic Labs Team. Performance and structural coverage of the latest, in-development AlphaFold model. (2023). <https://storage.googleapis.com/deepmind-media/DeepMind.com/Blog/a-glimpse-of-the-next-generation-of-alphafold/alphafold_latest_oct2023.pdf>.

67 Punjani, A., Rubinstein, J. L., Fleet, D. J. & Brubaker, M. A. cryoSPARC: algorithms for rapid unsupervised cryo-EM structure determination. Nat Methods 14, 290–296, doi:10.1038/nmeth.4169 (2017).

68 Scheres, S. H. RELION: implementation of a Bayesian approach to cryo-EM structure determination. J Struct Biol 180, 519–530, doi:10.1016/j.jsb.2012.09.006 (2012).

69 Pettersen, E. F. et al. UCSF Chimera--a visualization system for exploratory research and analysis. J Comput Chem 25, 1605–1612, doi:10.1002/jcc.20084 (2004).

70 Pettersen, E. F. et al. UCSF ChimeraX: Structure visualization for researchers, educators, and developers. Protein Sci 30, 70–82, doi:10.1002/pro.3943 (2021).

71 Liebschner, D. et al. Macromolecular structure determination using X-rays, neutrons and electrons: recent developments in Phenix. Acta Crystallogr D Struct Biol 75, 861–877, doi:10.1107/S2059798319011471 (2019).

72 Emsley, P., Lohkamp, B., Scott, W. G. & Cowtan, K. Features and development of Coot. Acta Crystallogr D Biol Crystallogr 66, 486–501, doi:10.1107/S0907444910007493 (2010).

73 Chen, V. B. et al. MolProbity: all-atom structure validation for macromolecular crystallography. Acta Crystallogr D Biol Crystallogr 66, 12–21, doi:10.1107/S0907444909042073 (2010).

